# A longevity-associated ubiquitin E3 ligase UBE3C governs lamin B1 homeostasis through selective autophagy and delay senescence

**DOI:** 10.64898/2026.08.05.741138

**Authors:** Di Guan, Seungsoo Kim, Karim Omar, Ying Hao, Gordon Huang, HyeRim Han, Jiping Yang, Naftali Horwitz, Phuong-Anh Dinh, Jihyun Hwang, Haiyuan Yu, Yousin Suh

**Author notes:** Corresponding author: Di Guan,; Yousin Suh,.

## Abstract

Nuclear lamina integrity is fundamental to cellular homeostasis across the lifespan ^1^, and its progressive deterioration is closely linked to human aging ^2^. Yet, the regulatory mechanism that govern this decline and how they might be counteracted in long-lived individuals remain poorly defined. Here, by combining whole-exome sequencing of Ashkenazi Jewish centenarians with GTEx transcriptomes, we identify ubiquitin E3 ligase UBE3C strongly associated with exceptional longevity and progressively declines with age across human tissues. UBE3C knockdown triggers premature senescence and destabilizes key nuclear lamina components Lamin B1 (LMNB1) and Lamin B receptor (LBR), while the longevity-associated UBE3C variant delays senescence and preserves LMNB1/LBR expression. Mechanistically, UBE3C interacts directly with LMNB1/LBR and modulates their abundance via selective autophagy. Notably, we uncover the ER-resident autophagy trigger CKAP4 ^3^ bridges UBE3C and LMNB1. UBE3C loss enhances LMNB1-CKAP4 binding, linking nuclear lamina turnover to autophagy. Together, our findings establish UBE3C as a central guardian of nuclear lamina maintenance during senescence and offering novel insights into interventions against age-related nuclear lamina deterioration.

## Main

The nuclear lamina, a highly conserved protein meshwork beneath the inner nuclear membrane ^4^, serves as a critical structural and functional scaffold for the nucleus. Composed primarily of A- and B-type lamins along with associated proteins, this dynamic structure not only maintains nuclear mechanical stability but also regulates essential processes including chromatin organization, DNA replication and repair, mechanotransduction, and transcription regulation ^5–11^. Given its central role in cellular homeostasis, dysfunction of the nuclear lamina is associated with diverse pathologies ranging from developmental disorders to age-related degeneration ^12–17^.

Lamin B1 (LMNB1), a major B-type lamin, is a well-established marker of cellular senescence ^18, 19^. LMNB1 with its binding partner, lamin B receptor (LBR), undergo progressive decline during aging across multiple human tissues ^18–22^, particularly in brain ^14, 23^. This decline is associated with nuclear envelope invaginations, heterochromatin loss, and the onset of senescence ^13, 17, 24^. While transcriptional downregulation contributes to reduced LMNB1 levels ^10, 18, 25^, the post-translational mechanisms regulating its stability, such as phosphorylation, acetylation, and ubiquitination ^26^, remain poorly characterized. Notably, autophagy-mediated LMNB1 degradation has been implicated in oncogene-induced senescence^27^, and emerging evidence suggests LMNB1/LBR may represent targets for senolytic interventions ^28, 29^, However, the precise regulatory mechanism of LMNB1/LBR in aging and its potential intersection with longevity remains unknown.

Here, we identify the E3 ubiquitin ligase UBE3C as a novel regulator of LMNB1 and LBR homeostasis. While UBE3C is implicated in protein degradation pathways through its ubiquitination activity ^30–34^, its direct link with aging has not been explored. Our study reveals that UBE3C is strongly associated with exceptional longevity and progressively declines with age across human tissues. Loss of UBE3C drives premature cellular senescence whereas a centenarian-enriched UBE3C variant delays senescence. We further demonstrated that LMNB1/LBR are the potential substrates of UBE3C during senescence. UBE3C maintains LMNB1/LBR levels through a non-canonical, autophagy-dependent mechanism, which engages the ER-resident autophagy trigger CKAP4. These findings reveal that UBE3C safeguard nuclear lamina integrity during senescence, offering a novel and therapeutically targetable pathway for aging interventions.

## UBE3C emerges as a top candidate gene associated with human longevity and aging

Longevity represents a robust model of successful aging, with centenarians providing unique insights into genetic factors that may delay aging and extend both healthspan and lifespan. Building on a previous genomic analysis of exceptional longevity, which identified 84,804 rare coding missense variants through whole-exome sequencing of 515 Ashkenazi Jewish centenarians and 496 controls ^35^, we focused this study on variants showing both genetic association with extended lifespan, and age-related gene expression changes. Given the central role of protein quality control in aging, we first compiled a list of 1799 genes involved in protein degradation pathways ^36–38^ (Supplementary Table 1), and then we examined their age-related expression patterns using GTEx transcriptomic data. This analysis identified 1571 protein degradation genes with significant age-dependent expression alterations (87% of 1,799 protein degradation genes; adjusted p value (adj.P) < 0.05; Supplementary Table 2). Integrative analysis with the top 100 longevity-associated variants highlighted five candidate genes at the intersection of longevity-associated variants and age-dependent expression changes (Fig. 1b). The E3 ubiquitin ligase UBE3C emerged as the leading candidate, based on both its rare variant association (rs146755594; p value=9.84E-04), and age-related downregulation in human tissues. These findings nominate UBE3C as a novel candidate of aging regulation and lifespan determination.

**Fig. 1:**
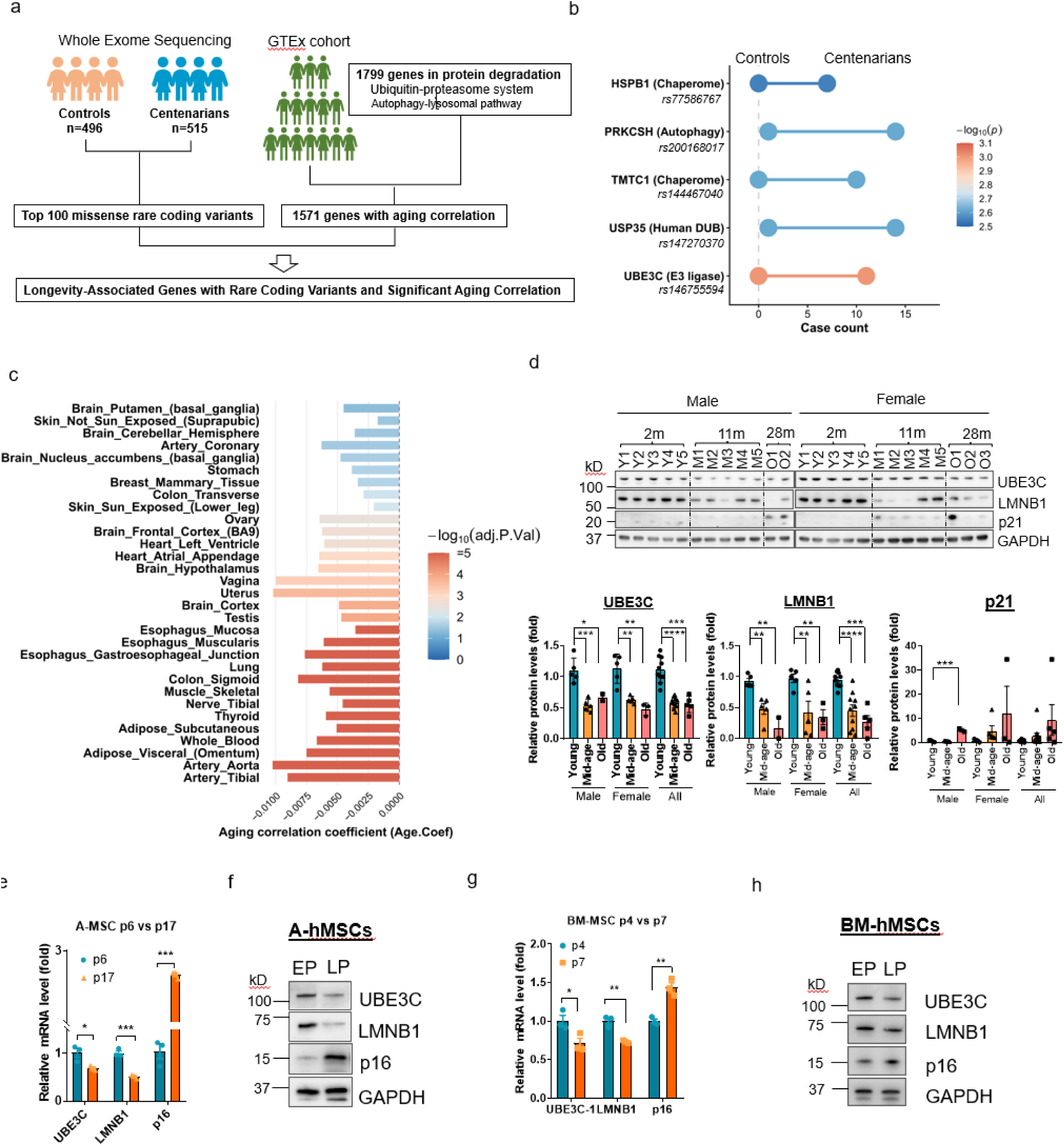
Ubiquitin E3 ligase UBE3C is highly associated with longevity and declines with age. a, Bioinformatics strategy for identifying longevity-associated rare coding variants with age-correlated expression. The whole-exome sequencing (WES) was performed on 496 controls and 515 centenarians to detect missense rare variants. Age-dependent transcriptomic analysis (GTEx) was applied to protein degradation pathways. Variants were prioritized by integration of top 100 missense rare coding variants with GTEx RNA-seq data, ranking by p value. **b**, Dumbbell plot showing counts of top 5 longevity-associated SNPs in controls (left) and centenarians (right). Gene names indicate the genes where these SNPs reside. Color intensity reflects the p value (−log_10_(p)) of SNP associations derived from WES data, highlighting genes strongly linked to longevity. SNP ID and annotation of genes were listed. **c,** Linear regression analysis of UBE3C expression (GTEx database) across human tissues revealed significant correlations with donor age (Adj.P.Val < 0.05), quantified by regression coefficients (Age.Coef). **d,** Western blot analysis (upper panel) of UBE3C, lamin B1 (LMNB1), and p21 in hippocampal lysates from C57BL/6 mice at 2 months (2m, young, n=10, male/female=5/5), 11 months (11m, middle-age, n=10, male/female=5/5), and 28 months (28m, old, n=5, male/female=2/3). Quantification (lower panel) is normalized to GAPDH. **e-h**, Analysis of UBE3C, LMNB1, and p16 (*CDKN2A*) levels in human mesenchymal stromal cells (hMSCs) at early (EP) and late passage (LP). qRT-PCR (**e**) and western blot (**f**) in adipose-derived MSCs (A- MSCs) at EP (p6, n=3) and LP (p17, n=3); parallel qPCR (**g**) and western blot (**h**) in bone marrow-derived MSCs (BM-MSCs) at EP (p4, n=3) and LP (p7, n=3). Error bars represent mean ± SEM; *p < 0.05, **p < 0.01, ***p < 0.001 (unpaired t-test unless noted).

## UBE3C exhibits an age-related decline

To investigate the relationship between UBE3C and aging, we first analyzed its expression patterns across a wide range of human tissues. Analysis of GTEx datasets revealed a significant age-related decline of UBE3C expression in 34 tissues, including artery (Tibial: adj.P=3.36E-26, aorta: adj.P=4.11E-13), subcutaneous adipose (adj.P=6.85E-10), skeletal muscle (adj.P=5.73E-08), and brain (adj.P=1.58E-4) (Fig. 1c and Extended Data Fig.1). Supporting these transcriptomic findings, human proteomics studies also showed a decrease in UBE3C protein abundance with age in human skeletal muscle and kidney ^39, 40^. This age-associated decrease in UBE3C is not unique to humans; analogous reductions in UBE3C mRNA and protein levels were observed in Drosophila during aging, underscoring the evolutionary conservation of this phenomenon ^41^. Consistently, immunoblotting of mouse hippocampus tissues confirmed a marked decline in UBE3C protein beginning at mid-age (11 months) and persisting into old age (28 months) compared to young controls (2 months) (Fig. 1d). Concordant changes in established aging markers, such as decreased Lamin B1 and increased p21, were also observed.

To assess UBE3C expression during cellular senescence-a fundamental hallmark of aging ^42^, we established replicative cellular senescence models using primary human mesenchymal stromal cells (hMSCs) from adipose (A-hMSCs) and bone marrow (BM-hMSCs). Both models recapitulated canonical senescence features upon serial passaging, including reduced LMNB1 and elevated p16 expression (Fig. 1e-h). Notably, we observed coordinated downregulation of UBE3C at both mRNA and protein levels in senescent cells. This *in vitro* pattern precisely mirrored the age-associated UBE3C decline observed *in vivo*, suggesting conserved regulation across cellular and organismal aging contexts.

## Depletion of UBE3C accelerates premature senescence

To investigate the contributions of UBE3C in cellular senescence, we stably knocked down UBE3C in hMSCs using two independent shRNAs, mimicking its age-dependent decline (Fig. 2a). UBE3C-knockdown (UBE3C-KD) hMSCs exhibited premature proliferation arrest during serial passaging, with significantly reduced population doubling rates compared to control cells (Fig. 2b). Cell cycle analysis revealed an accumulation of UBE3C-KD cells in G0/G1 phase and a corresponding decrease in S-phase cells (Fig. 2c), consistent with senescence-associated cell cycle arrest. This proliferative decline was accompanied by a marked increase in senescence-associated β-galactosidase (SA-β-gal)-positive cells (∼50% in UBE3C-KD vs ∼20% in controls; Fig. 2d). At the molecular level, UBE3C-KD hMSCs showed biomarkers of senescence, including reduced LMNB1 expression and elevated levels of the cell cycle inhibitors p16 and p21 (Fig.2e, f ). RNA-seq analysis revealed widespread transcriptomic changes, with 54 genes significantly upregulated and 108 genes downregulated in UBE3C-KD cells (adj.P < 0.05, |log2FoldChange|>1) (Extended Data Fig.2a). Gene ontology analysis of downregulated genes highlighted enrichment in chromatin organization and cell cycle regulation, (Fig. 2g), while upregulated genes were associated with extracellular matrix remodeling (Extended Data Fig. 2b), consistently with characteristic features of senescent cells.

**Fig. 2:**
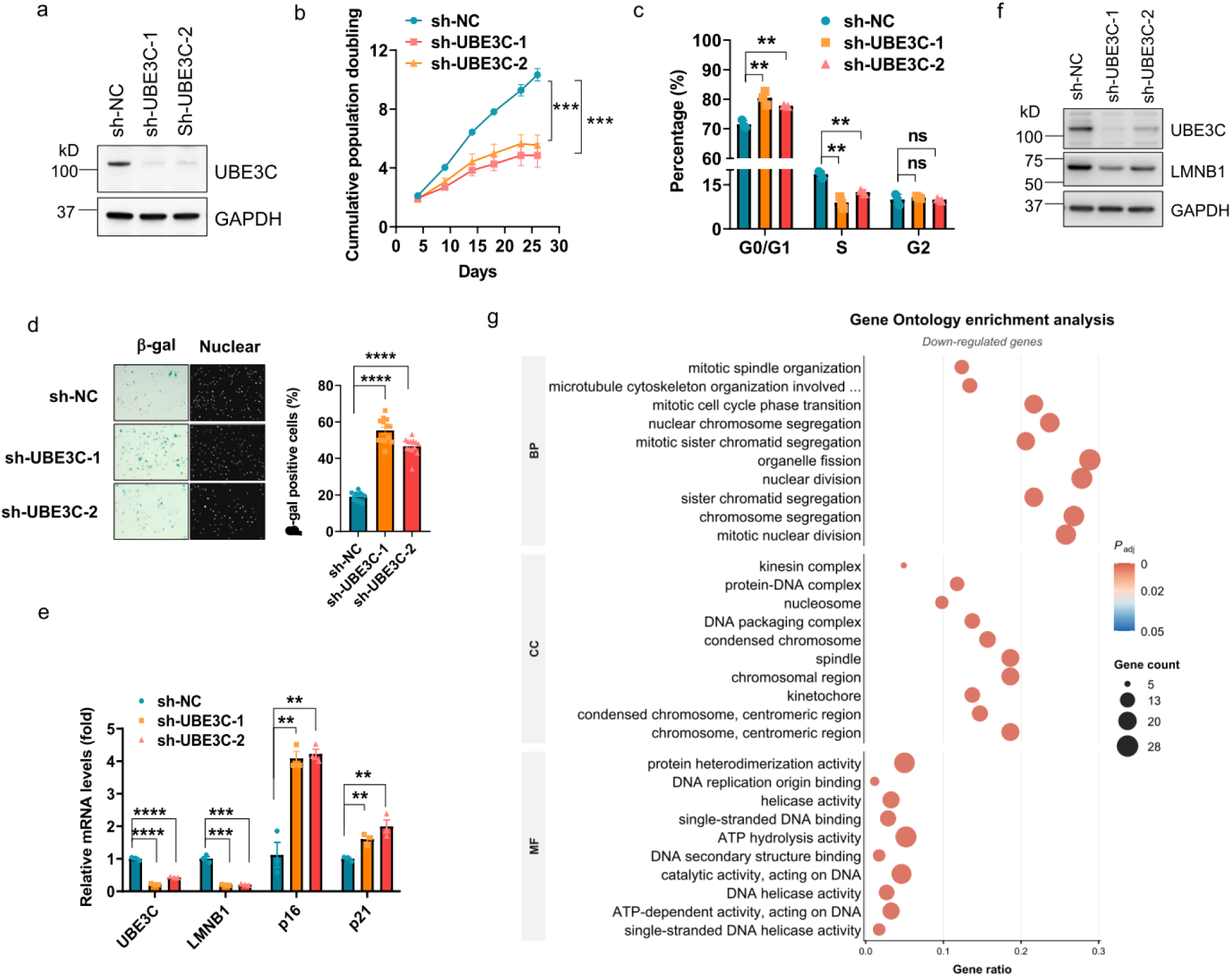
UBE3C depletion induces premature senescence in hMSCs. a, Western blot analysis of UBE3C protein levels in hMSCs transduced with lentiviral shRNAs targeting *UBE3C* (sh-UBE3C-1 and -2) or a non-targeting scramble control (sh-NC). Cells were analyzed 72 hours post-transduction and used for all subsequent panels. **b,** Population doubling curve of hMSCs following UBE3C knockdown. **c,** Cell cycle analysis by flow cytometry of hMSCs at passage 2 (day 8 post-transduction) and the percentage of cells in G0/G1, S, and G2 phases were quantified. **d,** Senescence-associated β-galactosidase (SA-β-gal) staining (left) and quantification of β-gal-positive cells (right) of hMSCs at passage 2 (day 8). Each dot represents an independent imaging field (n = 12 fields from 3 biological replicates). **e,** qRT- PCR analysis of *UBE3C*, *LMNB1*, *CDKN2A* (p16), and *CDKN1A* (p21) mRNA levels in hMSCs at passage 2 (day 8)**. f,** Western blot analysis of UBE3C and lamin B1 (LMNB1) protein expression in hMSCs at passage 2 (day 8)**. g,** Gene Ontology (GO) enrichment analysis of downregulated differentially expressed genes (DEGs; adj.P < 0.05, |log2FoldChange|>1; n=*2*) in hMSCs at passage 1 (day 4 post-transduction), covering biological process (BP), cellular component (CC), and molecular function (MF) categories. The top 10 pathways were shown. Error bars represent mean ± SEM. n=3 biologically independent experiments. *p < 0.05, **p < 0.01, ***p < 0.001 (unpaired t-test unless noted).

Collectively, these data demonstrate that UBE3C depletion drives hMSCs into premature senescence.

## Longevity-associated UBE3C variant extends cellular lifespan

To investigate UBE3C function in human longevity, we next examined the functional consequences of the longevity-associated rare coding variant in *UBE3C* gene (rs146755594 G→A) we prioritized. Initial characterization of the longevity-associated UBE3C variant (UBE3C^cent/cent^) through re-expression in human MSCs revealed a pronounced pro-longevity phenotype, including extended cellular lifespan (Fig. 3a) and reduced SA-β-gal positive senescence cells compared to UBE3C^+/+^ controls (Fig. 3b). To examine the variant’s effects in an endogenous context, we generated homozygous UBE3C^cent/cent^ knock-in human embryonic stem cells (hESCs) (Fig. 3c) and differentiated them into hMSCs. While UBE3C^cent/cent^ hESC-derived MSCs showed normal expression patterns of surface markers, they displayed remarkable resistance to replicative senescence (Fig. 3 d, e) and stress-induced senescence (Fig. 3f), evidenced by sustained proliferation (Fig. 3 d) and reduced SA-β-gal activity (Fig. 3e, f). UBE3C^cent/cent^ hMSCs maintained higher levels of LMNB1 and LBR (Fig. 3g), while UBE3C^cent/cent^ hESCs showed no differences in LMNB1 and LBR expression compared to wild type hESCs (Extended Data Fig.3). This stabilization of the nuclear lamina likely underlies the resilience conferred by the longevity-associated UBE3C variant, countering age-related senescence driven by lamina deterioration.

**Fig. 3:**
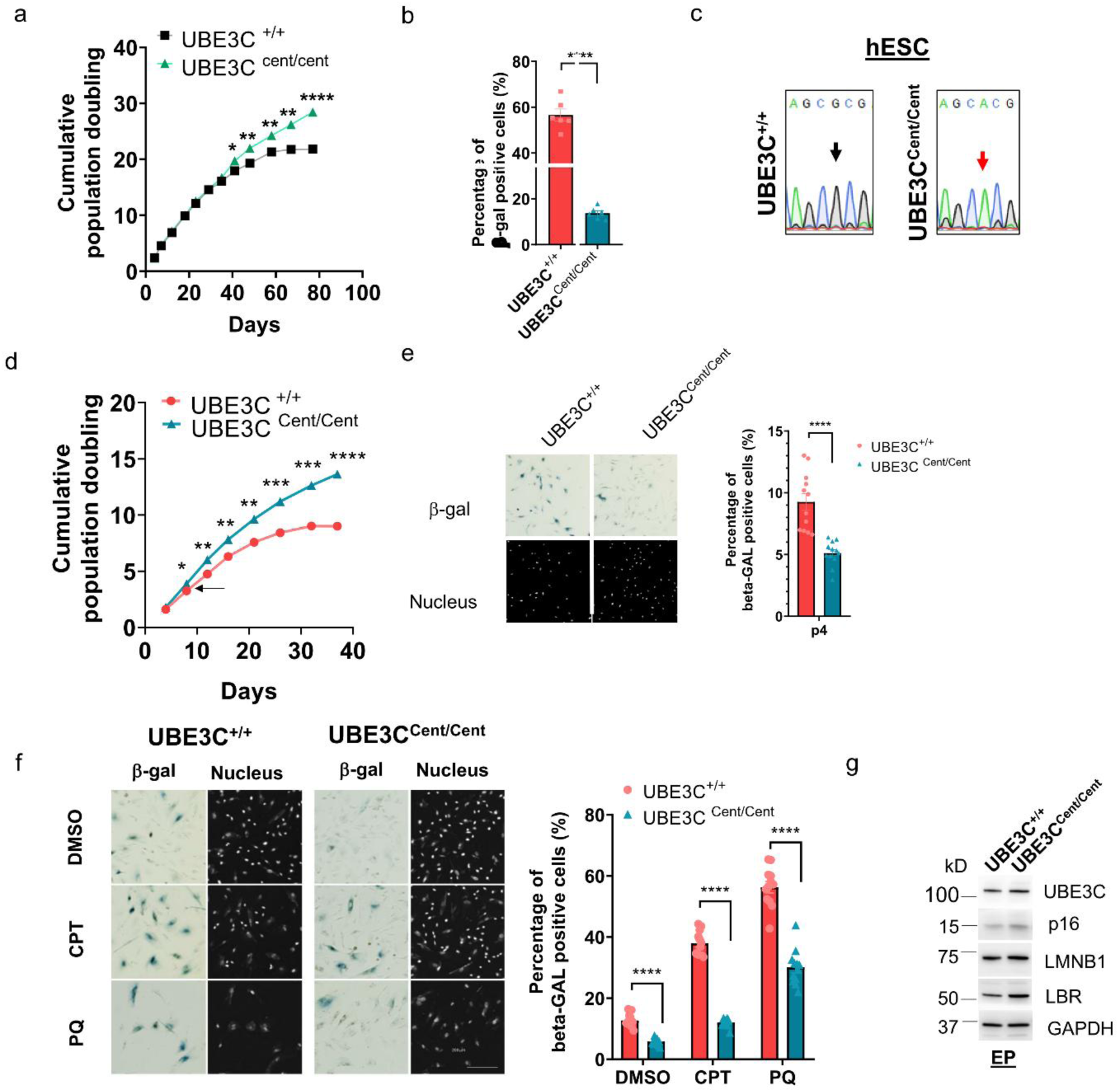
Longevity-associated UBE3C variant delays senescence and increases LMNB1/LBR in hMSCs. **a**, Population doubling curve of hMSCs transduced with lentivirus expressing either longevity-associated UBE3C variant (UBE3C*^cent/cent^*) or wild type UBE3C (UBE3C*^+/+^*). **b**, Quantification of β-gal-positive hMSCs transduced with lentivirus expressing either longevity-associated UBE3C variant (UBE3C*^cent/cent^*) or wild type UBE3C (UBE3C*^+/+^*) at passage 9 (day 48 post-transduction). Each dot represents an independent imaging field (n = 12 fields from 3 biological replicates). **c**, Sanger sequencing validation of the longevity-associated UBE3C variant in UBE3C*^cent/cent^* hESCs after CRSPR/cas9 knock-in. Chromatograms show the reference allele (G) and alternative allele (A) at the target locus in UBE3C*^cent/cent^* and UBE3C*^+/+^* hESCs. **d**, Population doubling curve of hMSCs derived from UBE3C*^cent/cent^* or UBE3C*^+/+^* hESCs, and **e**, SA-β-gal staining (left) and quantification (right) in hESCs-derived hMSCs at passage 4 (day 16). Each dot represents an independent imaging field (n = 12 fields from 3 biological replicates). **f**, SA- β-gal staining (left) and quantification (right) of hESC-derived hMSCs at passage 2 (day 8) following treatment with 2 μM cisplatin (CPT) or 1 mM paraquat (PQ) for 24 hours, followed by 48-hour recovery. DMSO served as a vehicle control. Each dot represents an independent imaging field (n = 12 fields from 3 biological replicates). **g**, Western blot analysis of UBE3C, LMNB1, LBR, and p16 (senescence marker) in hESC-derived hMSCs at passage 2 (day 8). GAPDH served as loading control. Error bars represent mean ± SEM (n = 3 biologically independent experiments unless noted). *p < 0.05, **p < 0.01, ***p < 0.001 (unpaired two- tailed Student’s t-test).

## UBE3C targets LMNB1 and LBR during senescence

To identify potential substrates of UBE3C during aging-related senescence, we combined quantitative proteomic profiling of UBE3C-KD and control hMSCs with UBE3C interactome analysis (Fig. 4a). The quantitative proteomic analysis revealed 48 upregulated and 105 downregulated proteins (adj.P<0.05, |log2FoldChange|>0.5) in UBE3C-KD cells compared to controls (Fig.4b and Extended Data Fig. 4a). The GO enrichment analysis showed that the downregulated proteins were significantly enriched for nuclear-localized factors involved in DNA/RNA metabolic processes and chromosomal organization (Extended Data Fig. 4b), while upregulated proteins predominantly participated in extracellular matrix remodeling (collagen metabolism), endocytosis, and cytoskeletal organization (Extended Data Fig. 4c). These proteomic changes showed remarkable concordance with our transcriptomic data from UBE3C- KD hMSCs (Fig. 2g and Extended Data Fig. 2b).

**Fig. 4:**
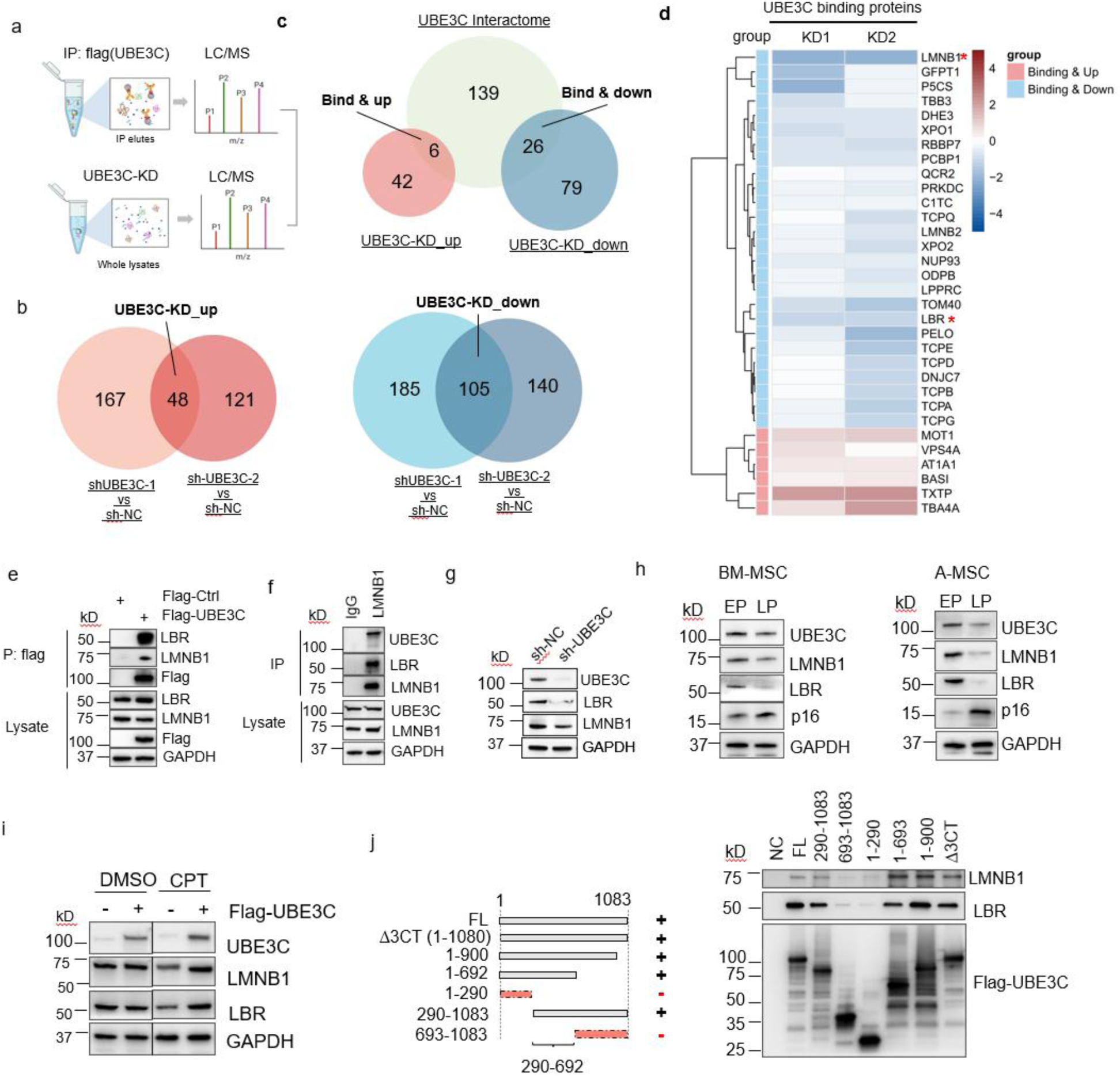
LBR/LMNB1 are substrates of UBE3C during cellular senescence. a–d, Integrated mass spectrometry (MS) analysis identifies potential UBE3C substrates. **a**, Schematic of the experimental workflow. Upper panel: HEK-293T cells expressing Flag-UBE3C or control were subjected to anti-Flag immunoprecipitation followed by LC/MS to define the UBE3C interactome. Lower panel: Quantitative LC/MS of human mesenchymal stem cells (hMSCs) transduced with two independent UBE3C shRNAs (UBE3C-KD; 72 h post-transduction) or control shRNAs identified differentially expressed proteins (DEPs) upon UBE3C depletion. **b**, Venn diagram shows overlapping DEPs between shUBE3C-1 and hUBE3C. Upregulated DEPs are shown in red, downregulated DEPs in blue. UBE3C-KD_up and UBE3C-KD_down indicate overlapping upregulated and downregulated DEPs, respectively. **c**, Venn diagram shows comparison between the UBE3C interactome and DEPs. The UBE3C interactome is indicated in blue. Binding & up: UBE3C binding protein upregulated upon UBE3C-KD; Binding & down: UBE3C binding proteins downregulated upon UBE3C-KD. **d**, Heatmap of the Binding & up and Binding & down proteins. LMNB1 and its bind receptor LBR (marked with red asterisk) are top candidates. **e**, co-IP validates UBE3C binding to LMNB1 and LBR in HEK-293T cells expressing flag-UBE3C or flag control. **f**, Endogenous co-IP using anti-LMNB1 antibody validates the binding of LMNB1 and LBR with UBE3C in hMSCs. **g**, Western blot analysis of UBE3C, lamin B1 (LMNB1), and lamin B receptor (LBR) in hMSCs at day 4 post-UBE3C knockdown. **h**, Western blot analysis of UBE3C, LMNB1, LBR, and p16 in BM-MSC (left) and A-MSC (right) at early passage (EP) and late passage (LP). **i** Western blot of UBE3C, LMNB1, and LBR in hMSCs expressing Flag-UBE3C or Flag control, treated with 1μM cisplatin (CPT) or DMSO for 24 h and followed by 48 h recovery. **j**, Protein interaction mapping defined the UBE3C 290-692aa domain as mediating LMNB1/LBR binding. Left panel: Schematic of UBE3C deletion constructs, including full-length (FL, 1-1083), catalytic-dead mutant (Δ3CT, 1-1080), catalytic domain deletions (1-900, 1-692), C-terminal fragments (290-1083, 693-1083), and empty vector control (NC). Dotted pink bars indicate constructs with impaired LMNB1/LBR binding. Right panel: Co-immunoprecipitation (Flag-IP) of LMNB1 and LBR from HEK-293T cells expressing UBE3C deletion mutants, analyzed by immunoblotting.

We next performed an interactome analysis to identify direct substrates of UBE3C. Immunoprecipitation-mass spectrometry (IP-MS) identified 171 high-confidence UBE3C-interacting proteins (Extended Data Fig. 5a), with significant enrichment in pathways involving protein folding and stability, ATP metabolic process, and cell cycle regulation (Extended Data Fig. 5b). Subcellular localization analysis showed these binding partners predominantly associate with microtubules and organelle membranes, including nuclear envelope (particularly nuclear lamina components), mitochondrial, and lysosomal compartments (Extended Data Fig. 5c). These findings link the function of UBE3C with nuclear lamina structure, mitochondrial homeostasis and lysosomal-mediated protein degradation.

By integrating the interactome of UBE3C with quantitative proteomic profiling of UBE3C-KD vs control hMSCs, we discovered 6 upregulated and 26 downregulated UBE3C-interacting proteins upon UBE3C knockdown (Fig. 4c, Extended Data Fig. 6). Interestingly, nuclear lamina components, particularly LMNB1 and LBR, emerged as top candidates among downregulated proteins in UBE3C-KD cells (Fig.4d, Extended Data Fig. 6). We confirmed these interactions through co-immunoprecipitation assays, demonstrating direct interaction between UBE3C and both LMNB1 and LBR (Figure 4e, f). Immunoblotting analysis showed marked reduction of LMNB1 and LBR protein levels upon UBE3C knockdown (Figure 4g).

LMNB1 and LBR are essential nuclear lamina components, whose dysfunction drives premature senescence and age-related tissue degeneration ^13, 14, 24, 29^. Our analysis of human GTEx data revealed significant age-associated declines of LMNB1 (Extended Data Fig. 7a) and LBR (Extended Data Fig. 7b) across multiple tissues. This age-associated decline was recapitulated in our UBE3C-deficient (Fig. 4g) and replicative senescent hMSC models (Fig. 4h), where both LMNB1 and LBR decline with loss of UBE3C. Furthermore, *In vivo* deletion of UBE3C showed decline of LMNB1 in UBE3C^-/-^ and UBE3C^+/-^ mice ear samples compared to the wild type control (Extended Data Fig. 8). Notably, complementation experiments demonstrated that UBE3C overexpression significantly attenuated the decline of both LMNB1 and LBR protein levels in DNA damage-induced senescent hMSCs (Fig. 4i). These findings suggest UBE3C as a guardian of LMNB1 and LBR protein stability during senescence.

## UBE3C interacts with LMNB1/LBR via 290-693 residues

To uncover the binding domain of UBE3C with LMNB1/LBR, we performed a domain mapping experiment using truncation mutants of UBE3C (Fig. 4j). While UBE3C fragments containing only the C-terminal catalytic domain (693-1080aa) or N-terminal (1-290aa) domains showed minimal binding to LMNB1/LBR, constructs maintaining the central region (1-1080aa, 290-1083aa, 1-900aa, or 1-692aa) preserved interaction capacity. These results demonstrate that the domain containing 290-692 residues constitutes the primary binding interface for UBE3C’s association with LMNB1/LBR, while the extreme N- and C-terminal catalytic regions are dispensable for this interaction.

## UBE3C protects LMNB1/LBR from autophagic degradation

To elucidate the mechanism underlying UBE3C-mediated LMNB1/LBR turnover, we first performed endogenous IP-MS to compare LMNB1 interactomes in UBE3C- depleted versus control hMSCs (Fig.5a). This revealed 15 differentially binding proteins (DBPs) of LMNB1, including 2 with enhanced and 13 with reduced binding upon UBE3C knockdown (adj.P < 0.05) (Fig. 5b-c, Extended Data Fig. 9). Interestingly, the DBPs ANXA2 and CKAP4 are key factors in autophagy ^3, 43, 44^, which showed increased binding with LMNB1 upon UBE3C loss (Fig. 5c, Extended Data Fig. 9). Cross-referencing the DBPs of LMNB1 with the UBE3C interactome identified 8 overlapping proteins, including CKAP4, PFKAL, PRKDC, LMNB2, TBA1B/TBA1C/TBB4B, and HS71B (Fig. 5d). Notably, the ER-resident autophagy trigger CKAP4 is the only up-regulated DBPs of LMNB1 and interacted with UBE3C, indicating CKAP4 bridges UBE3C with LMNB1 and links LMNB1 to autophagy degradation. Consistent with our findings, a previous study using lysosomal immunoprecipitation demonstrated that CKAP4 mediates LMNB1 recruitment to lysosomes, where UBE3C is detected ^45^. Altogether, our data suggests CKAP4 facilitates autophagy-lysosomal targeting of LMNB1 when UBE3C is deficient.

**Fig. 5.**
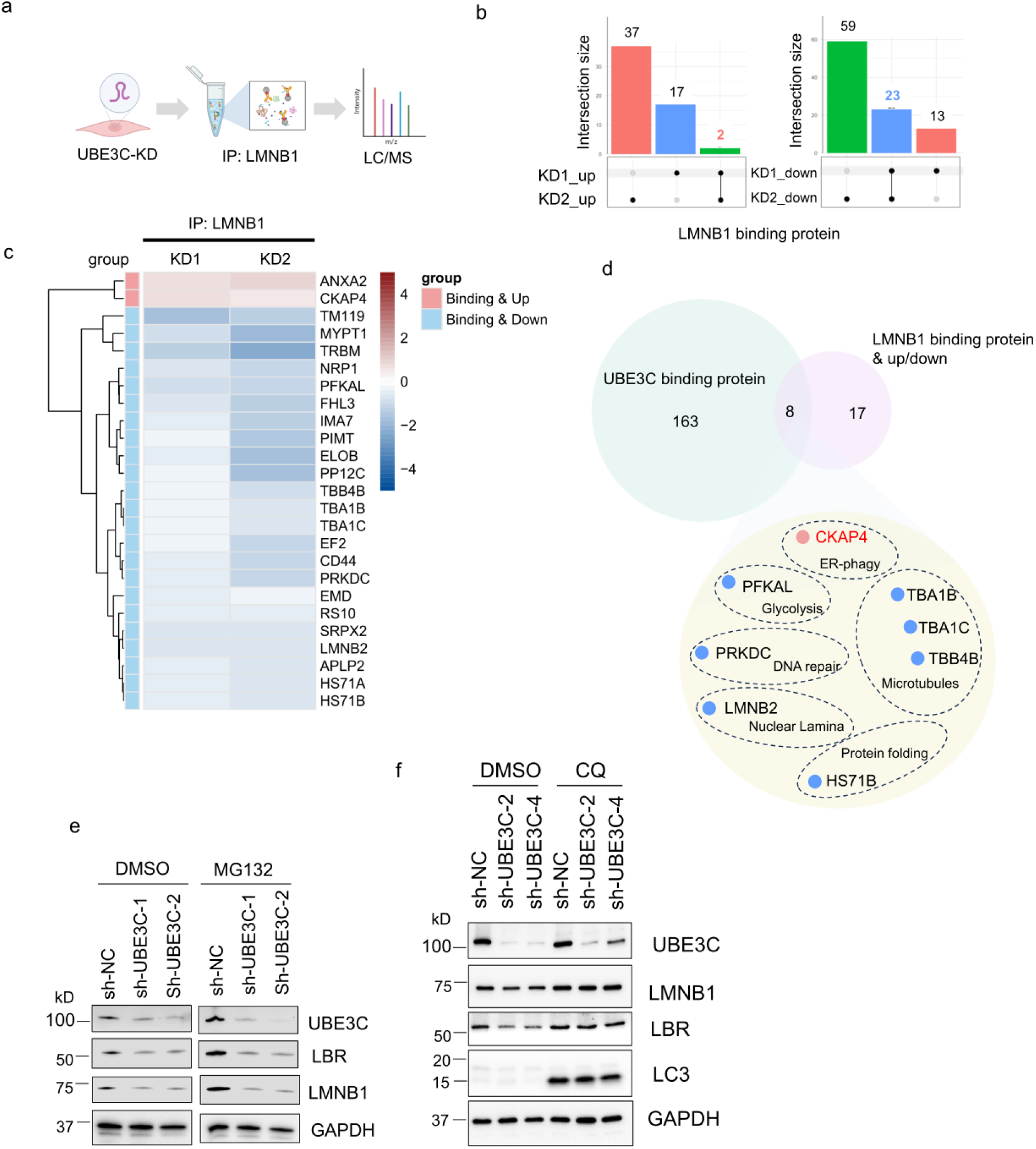
: UBE3C regulates Lamin B1/LBR via the nucleophagy pathway. a–d, Proteomic analysis of LMNB1 interactome alterations upon UBE3C depletion. **a**, Schematic of endogenous LMNB1 immunoprecipitation (IP) followed by LC/MS in hMSCs at passage 2 (day 10 post-transduction) with two independent shRNAs targeting UBE3C (UBE3C- KD) or non-targeting control. n = 3 biologically independent experiments. **b,** Upset plot shows overlapping differentially binding proteins (DEPs) of LMNB1 between two UBE3C-KD groups (KD1, KD2). Left panel: upregulated DBPs; Right panel: downregulated DBPs. Adj.p <0.05. **c**, Heatmap of upregulated (Binding & up) and downregulated (Binding &down) DBPs of LMNB1 upon UBE3C-KD as described in **b**. **d**, Venn diagram shows overlap between UBE3C-binding proteins (light blue) and DBPs of LMNB1 upon UBE3C depletion (pink). Overlapping proteins and their annotations are highlighted in light yellow. Proteins with increased LMNB1 binding are marked with pink dots; those with decreased binding are marked with blue dots. **e**, Western blot analysis of UBE3C, LMNB1, and LBR in hMSCs (3 days post-transduction) treated with 10 µM MG132 (proteasome inhibitor) or DMSO (6 hr). GAPDH served as loading control. **f**, Western blot analysis of UBE3C, LMNB1, and LBR in hMSCs (3 days post-transduction) treated with 50 µM chloroquine (CQ; lysosome inhibitor) or DMSO (24 hr).

To further confirm the autophagy-mediated degradation of LMNB1/LBR, we performed rescue assays on LMNB1 and LBR1 protein level by selectively inhibiting autophagy-lysosome pathways or the ubiquitin-proteasome system (UPS) in UBE3C- KD hMSCs. Treatment with the UPS inhibitor MG132 failed to restore Lamin B1/LBR levels in UBE3C-KD hMSCs (Fig. 5e). Additionally, UBE3C depletion did not alter LMNB1 ubiquitination levels but did alter the acetylation level (Extended Data Fig. 10). Together with our domain mapping result showing that UBE3C’s catalytic ubiquitin ligase domain is dispensable for LMNB1/LBR binding (Fig. 4i), our data supports a non-canonical mechanism of UBE3C on LMNB1 regulation. In contrast, the autophagy inhibitor chloroquine (CQ) rescued LMNB1/LBR levels in UBE3C-KD cells, at least partially (Fig. 5f). Together, these findings establish UBE3C as a gatekeeper of nuclear lamina homeostasis that functions through a non-canonical, catalysis-independent mechanism to prevent autophagic degradation of LMNB1/LBR during senescence (Fig. 4g).

In summary, our study identifies UBE3C as a novel and critical regulator of LMNB1 and LBR during aging-related senescence. Integrative analysis of centenarian whole-exome sequencing and GTEx transcriptomes pinpoints UBE3C as a top longevity-associated gene that declines with age. UBE3C loss drives premature cellular senescence and destabilization of LMNB1/KBR, while the longevity-associated UBE3C variant preserves LMNB1/LBR levels and confers resilience. By integrating proteomic and functional analyses, we reveal a non-canonical quality control mechanism in which UBE3C selectively protects LMNB1 and LBR from autophagic degradation. These findings define the UBE3C-LMNB1/LBR axis as a key determinant of senescence cell fate, and link LMNB1/LBR turnover to autophagy machinery, providing potential therapeutic entry points for mitigating aging-related nuclear lamina deterioration.

## Discussion

In this study, we identified UBE3C as a novel longevity-associated E3 ligase that preserves nuclear lamina integrity to counteract cellular senescence. By integrating whole-exome sequencing of Ashkenazi Jewish centenarians with age-stratified GTEx transcriptomes, we pinpointed UBE3C as a top candidate that exhibits both genetic association with exceptional longevity and progressive age-related decline across human tissues. Functional assays confirmed that UBE3C depletion triggers premature senescence, while the longevity-associated UBE3C variant confers resilience. Through integrated proteomic and interaction analyses, we demonstrated that UBE3C interacts with the nuclear lamina components LMNB1/LBR and prevents their autophagic degradation during senescence. These findings establish UBE3C as a key regulator of nuclear lamina integrity and uncover a non-canonical, autophagy-linked quality control mechanism.

While UBE3C is established as an E3 ubiquitin ligase involved in protein degradation ^30–34^, we uncovered a UPS-independent mechanism for its regulation on LMNB1/LBR. Instead, UBE3C prevents their autophagic turnover, a finding that expands the functional repertoire of ubiquitin ligases in aging. Notably, this occurs through an LC3-independent pathway (Extended Data Fig. 11), distinguishing it from autophagy-mediated LMNB1 degradation during oncogenic stress-induced senescence ^27^. Our interactome data implicates the ER resident autophagy triggers CKAP4 in this process, consistent with prior reports of their colocalization in lysosomes. These results suggest a model where UBE3C functions as a guardian that prevents CKAP4-mediated lysosomal targeting of the nuclear lamina, although further validation is required.

As a well-established hallmark of aging ^42^, cellular senescence drives tissue dysfunction and age-related pathologies ^46^. Our work positions UBE3C as a key regulator of senescence through its stabilization of LMNB1/LBR via autophagy-lysosomal inhibition. This mechanism offers new targets for aging-related senescence intervention strategies.

Longevity genetics provides a powerful lens to identify protective aging pathways ^47^. While many longevity-associated variants have been reported, few have been mechanistically dissected. Here, using isogenic knock-in models and functional assays, we demonstrated that a centenarian-enriched UBE3C variant (rs146755594) delays both replicative and stress-induced senescence by maintaining LMNB1/LBR levels. Potential future *in vivo* studies can further validate its pro-longevity effects.

In conclusion, our study uncovers UBE3C as a key longevity-associated regulator that preserves nuclear lamina integrity by stabilizing LMNB1 and LBR through an autophagy-lysosomal mechanism, independent of the UPS. These findings establish a novel proteostatic pathway linking UBE3C-mediated nuclear lamina maintenance to both cellular senescence and longevity. While our current work primarily demonstrates this mechanism *in vitro*, future investigations employing *in vivo* models and diverse cell types will be critical for translating these discoveries into therapeutic strategies. This study not only advances our fundamental understanding of nuclear lamina homeostasis in aging but also provides a foundation for developing targeted interventions to improve healthspan.

## Supporting information

Genes in protein degradation pathways

Protein degradation genes with significant aging correlation in human tissues

Tissue sample sizes in GETx analysis

Antibodies used in this study

Primers for qRT-PCR

## Acknowledgement

We acknowledge the Columbia Genome Center for bulk RNA sequencing, and the flow cytometry core for FACS sorting. We thank the members of the Suh lab for discussions and supports, and our former technician Ashley Wilczek for help on the plasmid constructions. We appreciate the plasmids kindly shared from Zhixun Dou lab. This work is supported by National Institutes of Health (AG069750, DK127778, AG057433, AG061521, HL150521, AG055501, AG057341, AG057433, AG057706, AG057909 and

AG017242 to Y.S.), a grant from the Simons Foundation (to Y.S.); and the Impetus Grant Program (to Y.S.).

## Contributions

D.G. and Y.S. conceived and designed the study. D.G. performed most of the experiments, analyzed and interpreted the data, and wrote the manuscript with the output of all authors. S.K. performed the GTEx analysis and provided analytic advice for WES and RNA-seq. K.O. conducted the quantitative MS on UBE3C-KD versus control hMSCs and UBE3C interactome under the supervision of H.Y. Y.H. contributed part of the quantitative MS analysis. G.H and H.H. assisted with IP and WB experiments.

J.Y. provided analytic advice on WES and RNA-seq, and assisted with genetic editing in hESCs. J.H assisted with GTEx analysis. N.H. and A.D. reviewed the manuscript and provided advices. Y.S. reviewed the manuscript, oversaw the project, and secured funding.

## Methods

### Cell culture

Primary human adipose-derived mesenchymal stem cells (A-hMSC) and bone marrow-derived mesenchymal stem cells (BM-hMSC) were cultured in α-MEM (Gibco) containing 10% FBS (Gibco), 1% penicillin-streptomycin (Gibco), and 1 ng/mL recombinant human bFGF (STEMCELL Technologies) on 0.1% gelatin (Invitrogen) -coated surfaces. H7 human embryonic stem cells (hESCs, WiCell WA07) were maintained in a feeder-free maintenance medium mTESR plus (STEMCELL technology) on matrigel (Corning)-coated plates. For differentiation, H7 were maintained on IR–treated CF1 mouse embryonic fibroblasts (MEF, Gibco) feeder in DMEM/F-12 medium (Gibco) with 20% knockout serum replacement (Gibco), 0.1 mM non-essential amino acids (Gibco), 55 μM β-mercaptoethanol (Thermo Fisher Scientific) and 10 ng ml−1 basic fibroblast growth factor (bFGF; STEMCELL Technologies). HEK293T cells were grown in DMEM (Gibco) supplemented with 10% FBS (GeminiBio) and 1% penicillin/streptomycin (Gibco). All cell lines were incubated at 37°C in a 5% CO₂ humidified incubator and regularly tested for mycoplasma contamination.

### GTEx analysis

The GTEx V8 RNA-seq profiles from all tissues ^43^ were analyzed to identify genes associated with age. We removed the tissues with a sample size of ≤ 100 (minimum sample size ≥ 164; Supplementary Table 3). For each tissue separately in the combined, male, and female cohorts, the associations between gene expression and age were tested using a linear regression analysis with the limma package (version 3.48.3). Nominal P-values were corrected for multiple testing by the Benjamini-Hochberg procedure, controlling the False Discovery Rate (FDR), and significant genes were selected by adjusted *P* < 0.05 and across ≥ 5 tissues.

### Prioritization of the top longevity-associated genetic candidates

The top longevity-associated genetic candidates were prioritized through an integrative three-step analysis. Initially, the top 100 rare coding variants associated with longevity were selected from a previously published whole-exome sequencing data of 515 Ashkenazi Jewish centenarians and 496 population-matched controls ^29^. Subsequently, protein degradation genes involved in ubiquitin-proteasome and autophagy-lysosomal pathways were compiled from published studies (Supplementary Table 1) ^40–43^. Age-correlation analysis of the genes was performed using GTEx v8 RNA-seq data, and significant genes were selected by adjusted p<0.05, across ≥5 tissues as described above. Intersection of the two candidate lists revealed the top 5 longevity-associated genetic candidates containing both: 1) previously reported longevity-associated rare coding variants and 2) significant age-correlated gene expression changes. The gene ranking was determined by association p-values of the centenarian-enriched rare coding variants from the original study.

### CRISPR/Cas9-mediated knock-in in human ESCs

The SNP rs146755594 was introduced into H7 embryonic stem cells using a CRISPR/Cas9 genome editing approach. Target-specific sgRNA and single-stranded donor oligonucleotide (ssODN) were designed using the IDT (the Integrated DNA Technologies) Alt-R CRISPR design platform. H7 ESCs were co-transfected with pCas9-GFP (Addgene #44719) expressing Cas9 nuclease, pCAGmcherry-gRNA plasmid targeting the rs146755594 locus, and ssODN donor template using Lipofectamine STEM transfection reagent according to the manufacturer’s protocol (Invitrogen). sgRNA: TGAGTCGTGACAGCTCGCGC. ssODN: G*G*CTTCTTCACTCTCCTCCTCAGAGTCACTGGCTGAGTCGTGACAGCTCGTGCTGGCAGGAGAGA CTGGTAACTGAGAGAGGAAGGTCTGCAGCACCCG*C*A (*phosphorothioate linkages).

At 48 hours post-transfection, enzymatically dissociated single cells (Accutase, STEMCELL Technologies) were sorted by FACS based on GFP and mCherry dual fluorescence. For clonal isolation, 2×10⁴ sorted cells were seeded onto a CytoSort™ microwell array (20,000 microwells, CELL Microsystems) pre-coated with Matrigel (Corning). Following 3 days of culture, individual colonies were automatically selected and transferred to 96-well plates using the CellRaft AIR System (CELL Microsystems). Expanded clones were screened using a custom TaqMan SNP Genotyping Assay (ThermoFisher) targeting rs146755594, with positive clones subsequently validated by Sanger sequencing of the edited locus.

### Derivation of hMSCs from hESCs

hMSCs were differentiated from H7 hESCs using a modified embryoid body-based protocol described previously ^40^. Briefly, hESC colonies were detached using dispase (0.5mg/ml) and transferred to low-attachment plates to permit embryoid body formation. The resulting embryoid bodies were subsequently plated onto Matrigel-coated 6-well plates and maintained in αMEM (Gibco) medium supplemented with 10% FBS (Gibco), 10 ng/ml bFGF (STEMCELL Technologies), 5 ng/ml TGFβ (STEMCELL Technologies) and 1% penicillin/streptomycin (Gibco). Following 14-21 days of differentiation, fibroblast-like cells emerging from the embryoid bodies were isolated by fluorescence-activated cell sorting (FACS) using a defined surface marker profile (CD73+/CD90+/CD105+). The purified hMSC population was expanded for subsequent experiments

### Growth curve assay

Cell proliferative capacity was quantitatively evaluated by cell population doubling numbers through serial passage monitoring following established protocols ^41^. In brief, proliferative capacity per passage was determined by applying the equation:

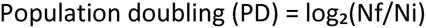

where Ni represents the initial seeded cell count and Nf denotes the final harvested cell number. The summation of PD values across successive passages generated cumulative population doubling curves.

### Cell cycle analysis

Cell cycle distribution was quantified using the Click-iT™ EdU Alexa Fluor™ 647 Flow Cytometry Assay Kit (Thermo Fisher Scientific) following the manufacturer’s protocol. In brief, Proliferating cells were pulse-labeled with 10 µM 5-ethynyl-2’-deoxyuridine (EdU) for 2 hours in complete medium before trypsinization (0.25% trypsin-EDTA, Gibco). Single-cell suspensions were simultaneously processed for EdU detection via Alexa Fluor™ 647 conjugation and DNA content staining with 5 µg/mL Hoechst 33342 (Thermo Fisher Scientific) for total DNA content. Samples were acquired on a BD LSRFortess cell analyzer, with subsequent cell cycle phase distribution analysis conducted using FlowJo v10.10 software.

### Plasmids and viral constructs

For UBE3C expression studies, the pcDNA3.1-Flag-UBE3C construct (GenScript) was used to express Flag-tagged UBE3C. Derivative constructs, including a Flag empty vector control (pcDNA3.1-Flag-Ctrl) and a C-terminal truncated mutant (pcDNA3.1-Flag-UBE3C-Δ3CT), were generated from the pcDNA3.1-Flag-UBE3C using the Q5® Site-Directed Mutagenesis Kit (New England Biolabs), according to the manufacturer’s instructions. For LMNB1 overexpression, the pcDNA3.1-6xHis-LMNB1 construct was generated by subcloning LMNB1 cDNA into RGS- 6xHis-pcDNA3.1 (Addgene) using the In-Fusion® Snap Assembly Master Mix (Takara). The pRK5-HA-Ubiquitin-WT plasmid (Addgene) was employed in ubiquitination assays. For LC3 or GABARAP expression, the Flag-LC3 and Flag-GABARAP plasmids were kindly shared by Dr. Zhixun Dou (Massachusetts General Hospital and Harvard University). Lentiviral constructs, including plenti-EGFP-UBE3C^+/+^ and plenti-EGFP-UBE3C^cent/cent^, were engineered from the plenti-cmv-msv-puro-EGFP backbone (Addgene) to express either the wild-type or a longevity-associated UBE3C variant. For gene silencing, LV10N-based lentiviral shRNA plasmids targeting UBE3C (LV10N-shUBE3C-1 and LV10N-shUBE3C-2) and a non-targeting control (LV10N-shNC) were designed and synthesized by GenePharma. Targeting sequence of sh- UBE3C-1: GCACTACATGATTCACAATGG, sh-UBE3C-2: GCAGACCATCCTGTTATTAAG, and sh-NC: GTTCTCCGAACGTGTCACGT. Lentiviral particle production was performed using psPAX2 and pMD2.G helper plasmids (Addgene).

### Lentiviral preparation and cell transduction

To generate lentiviral particles expressing either wild-type or the longevity-associated UBE3C variant, HEK293T cells were co-transfected with plenti-EGFP-UBE3C+/+ , plenti-EGFP- UBE3Ccent/cent, or an empty control, along with psPAX2 and pMD2.G packaging plasmids, using polyethylenimine (PEI; Sigma). For UBE3C knockdown, LV10N-shUBE3C-1, LV10N- shUBE3C-2, or a non-targeting control (LV10N-shNC) were similarly co-transfected with psPAX2 and pMD2.G using PEI. Viral supernatants were collected 48 hours post-transfection and concentrated using the Lenti-X™ Concentrator (Takara Bio) in accordance with the manufacturer’s protocol. Target human cell lines were transduced with the concentrated lentivirus in the presence of 4 µg/ml polybrene (Sigma). Following transduction, cells were selected with 1 ng/ml puromycin for 48 hours to establish stable populations for downstream analyses.

### Immunoprecipitation (IP)

For exogenous Co-IP assays, HEK293T cells were transfected with the indicated plasmids using polyethylenimine (PEI) and processed based on previously described protocols with modifications ^36,38^. In brief, cells were harvested and lysed in ice-cold IP buffer composed of 25 mM Tris-HCl (pH 7.4), 150 mM NaCl, 1 mM EDTA, 1% Triton X-100, and 5% glycerol, supplemented with 1× Pierce™ Protease and Phosphatase Inhibitor Cocktail (Thermo Fisher Scientific). Lysates were gently rotated at 4 °C for 30 minutes and cleared by centrifugation at 13,000 rpm for 15 minutes at 4 °C. The resulting supernatant was transferred to fresh tubes and incubated with the appropriate antibody-conjugated beads overnight at 4 °C under constant rotation. Beads were washed five times with IP buffer and bound proteins were eluted using Elution Buffer (100 mM Tris-HCl, pH 7.5, 1% SDS) by heating at 65 °C for 15 minutes. Eluted samples were analyzed by western blotting.

For endogenous Co-IP experiments, cells were lysed in a modified buffer (20 mM Tris-HCl, pH 7.5, 137 mM NaCl, 1 mM MgCl₂, 1 mM CaCl₂, 1% NP-40, and 10% glycerol) supplemented with Pierce™ Protease and Phosphatase Inhibitor Cocktail and 12.5 U/mL benzonase (Novagen) to enhance solubilization of chromatin-bound proteins ^21^. Lysates were rotated at 4 °C for 30–60 minutes, clarified by centrifugation at 13,000 rpm, and processed for immunoprecipitation as described above.

The following antibodies and beads were used: ANTI-FLAG® M2 Affinity Gel (Sigma), Anti-Lamin B1 antibody [EPR22165-121] (Abcam), Pierce™ Protein A/G Plus Agarose (Thermo Fisher Scientific).

### Ubiquitination Assay

To detect ubiquitinated proteins, we followed an established immunoprecipitation-based approach ^42^. After Co-IP, ubiquitinated species were detected by immunoblotting using anti-HA (C29F4) (CST) or anti-ubiquitin [EPR8830] antibodies (Abcam)

### Western blotting

Western blotting was performed as previously described ^40^. For cultured cells and IP elutes, lysates were prepared using Pierce™ Lane Marker Reducing Sample Buffer (Thermo Fisher Scientific) and denatured at 95°C for 5–10 minutes. Proteins were separated by SDS-PAGE and transferred to membranes for immunoblotting. Signals were visualized using the ChemiDoc Imaging System (Bio-Rad).

For mouse tissues, samples were lysed in ice-cold RIPA buffer (Thermo Fisher Scientific) on ice for 30 minutes, followed by brief homogenization and sonication. Lysates were clarified by centrifugation, and protein concentrations were measured using the Pierce™ BCA Protein Assay Kit (Thermo Fisher Scientific). Equal amounts of protein were mixed with Lane Marker Reducing Sample Buffer and processed for western blotting as described above.

A detailed list of the antibodies used is provided in Supplementary Table 4.

### Senescence-associated β-galactosidase (SA-β-GAL) staining

SA-β-GAL staining was performed as previously described ^40^. Briefly, cells were washed with PBS and fixed at room temperature for 2-5 minutes using a fixation buffer containing 2% formaldehyde and 0.2% glutaraldehyde. After fixation, cells were incubated overnight at 37 °C with freshly prepared staining solution. Images were acquired using the EVOS Cell Imaging System (Thermo Fisher Scientific). SA-β-GAL-positive and total cell numbers were quantified using ImageJ software, and the percentage of SA-β-GAL-positive cells was calculated and analyzed statistically using GraphPad Prime 8.

### RT-qPCR (quantitative reverse transcription polymerase chain reaction)

Total RNA was isolated using the RNeasy Plus Kit (Qiagen) according to the manufacturer’s instructions. One microgram of total RNA was reverse transcribed into complementary DNA (cDNA) using the PrimeScript™ RT Reagent Kit (Takara). Quantitative real-time PCR (qPCR) was conducted on a QuantStudio™ 6 Pro Real-Time PCR System (Applied Biosystems) using PowerUp™ SYBR™ Green Master Mix (Applied Biosystems). Gene expression levels were normalized to GAPDH, and relative expression was calculated using the ΔΔCq method. Primer sequences used for all RT-qPCR assays are provided in in Supplementary Table 5.

### Bulk RNA-sequencing

Total RNA was extracted from 1 × 10⁶ cells (per duplicate) using the RNeasy Plus Mini Kit (Qiagen) according to the manufacturer’s instructions. RNA integrity was assessed using the Agilent Bioanalyzer (Agilent Technologies). RNA libraries were constructed using the Ribo-Zero Plus rRNA Depletion Kit (Illumina), following the manufacturer’s protocol. The sequencing was performed on an Illumina NovaSeq 6000 platform to generate paired-end reads.

Sequencing reads were aligned to the human reference genome (hg38) using STAR aligner (v2.7.10a) and gene-level quantification was performed using featureCounts from the Subread package (v2.0.3). Differential gene expression analysis was conducted using DESeq2 (v1.38.1). Gene set enrichment analysis (GSEA) was performed using the clusterProfiler package (v4.4.1) in R. Volcano plots were generated using the ggplot2 package (v4.0.3) in R.

### Animal experiments

All animal experiments were approved by the Columbia University Irving Medical Center Institutional Animal Care and Use Committee (IACUC).

For hippocampal tissue collection, 4-week-old C57BL/6 mice were obtained from Jackson Laboratory and bred in-house. Hippocampi were harvested from mice at 2 months (n = 5 males, 5 females), 11 months (n = 5 males, 5 females), and 28 months (n = 2 males, 3 females) following euthanasia, and processed for western blotting.

For ear tissue collection, cryopreserved heterozygous UBE3C-null (UBE3C^+/−^) mice were purchased from Taconic Biosciences and revived in-house. These mice were backcrossed with C57BL/6 mice (Jackson Laboratory) and genotyped by Transnetyx. UBE3C^-/-^, UBE3C^+/-^, and UBE3C^+/+^ littermates were generated from UBE3C^+/-^ breeding pairs and genotyped by Transnetyx. Ear punches were collected from 3-month-old mice and processed for western blot analysis.

### Proteomics sample preparation

Sample preparation for DIA proteomics was conducted on a fully automated workflow as previously described ^48^. Briefly, 200 µL lysis buffer (50 mM Tris-HCl, 50 mM NaCl, 1% SDS, 1% Triton X-100, 1% NP-40, 1% Tween-20, 1% glycerol, 1% sodium deoxycholate (wt/vol), 5 mM EDTA, 5 mM dithiothreitol (DTT), 5 KU benzonase, and 1× complete protease inhibitor) was added to the samples. The lysates were incubated at 65 °C for 30 min at 1,200 rpm for protein denaturation. Samples were then alkylated with 10 mM iodoacetamide for 30 min at room temperature. Protein concentration was determined using the DC Protein Assay. Protein enrichment and on-bead tryptic/Lys-C digestion were performed using a KingFisher APEX robotic system. Proteins were digested with Trypsin/Lys-C mix in 50 mM ammonium bicarbonate at 37 °C for 18 h. After digestion, peptides were vacuum-dried and reconstituted in 2% acetonitrile (ACN) with 0.1% formic acid. A total of 1 µg of peptides was used for LC-MS/MS DIA analysis.

For TMT labeling, 100 µg of peptides from each sample were labeled with TMTpro 16-plex reagents (Thermo Fisher Scientific) according to the manufacturer’s instructions. Briefly, TMT reagents were equilibrated to room temperature and dissolved in anhydrous ACN. Peptides were reconstituted in 100 µL of 100 mM HEPES (pH 8.5) and mixed with TMT reagents at a 1:4 (peptide:TMT) weight ratio. After incubation for 1 h at room temperature, the reaction was quenched with 5% hydroxylamine for 15 min. Labeled samples were pooled at a 1:1 ratio, vacuum-dried, and desalted using C18 spin columns (Thermo Fisher Scientific). Desalted pooled peptides were then fractionated into 8–12 fractions using high-pH reversed-phase chromatography (Thermo Fisher Scientific). Each fraction was vacuum-dried and reconstituted in 2% ACN with 0.1% formic acid for LC-MS/MS analysis. A total of 1 µg of peptides from each fraction was used for LC-MS/MS analysis.

### Liquid chromatography and mass spectrometry (LC/MS)

For each DIA sample, digested peptides were analyzed using an UltiMate 3000 nano-HPLC system (Thermo Fisher Scientific) coupled with an Orbitrap Eclipse mass spectrometer (Thermo Fisher Scientific). Peptides were loaded at 5 µL/min onto a trap column (75 µm × 2 cm, PepMap nanoViper C18 column, 3 µm, 100 Å, Thermo Fisher Scientific) equilibrated in 2% ACN with 0.1% trifluoroacetic acid (TFA). Samples were loaded onto the trap column for 5 min and then switched in-line with an ES903A nano C18 column (75 µm × 500 mm, 2 µm, 100 Å, Thermo Fisher Scientific) using an 80-minute linear gradient of 2–40% phase B (5% DMSO in 0.1% formic acid in ACN). Column temperature was maintained at 60 °C and the flow rate was set to 300 nL/min. Data were acquired in data-independent acquisition (DIA) mode. The MS1 scan was set at a resolution of 120,000, with a standard AGC target, and the maximum injection time was set to auto. The MS2 scans covered a precursor mass range of 400–1,000 m/z, using an isolation window of 8 m/z with 1 m/z overlap, resulting in a total of 75 windows per scan cycle. Fragmentation was performed using high-energy collisional dissociation (HCD) with a normalized collision energy of 30%. MS2 spectra were acquired at a resolution of 30,000 with a scan range of 145–1,450 m/z, and the loop control was set to 3 seconds.

For TMT-labeled samples, LC-MS/MS analysis was performed using the same nano-HPLC system coupled with an Orbitrap Eclipse mass spectrometer. Peptides were loaded and separated as described above with the following modifications. Data were acquired in data-dependent acquisition (DDA) mode. MS1 scans were acquired at a resolution of 120,000 over a mass range of 400–1,500 m/z, with a standard AGC target and an automatic maximum injection time. The top 15 most abundant precursors were selected for MS2 fragmentation using HCD with a normalized collision energy of 35%. MS2 scans were acquired at a resolution of 50,000 with an isolation window of 0.7 m/z. A dynamic exclusion of 30 seconds was applied to avoid repeated sequencing of the same precursor. TMT reporter ions were detected in the MS2 spectra with the synchronous precursor selection (SPS) mode enabled (SPS = 10).

### Proteomics data analysis

DIA database searches were performed using Spectronaut (version 20, Biognosys) with the directDIA search strategy. The UniProt human proteome reference (reviewed genes, 20,384 entries) was used as the library FASTA file. The directDIA false discovery rate (FDR) was set to 1%. Protein N-terminal acetylation and methionine oxidation were set as variable modifications, and carbamidomethylation of cysteine residues was set as a fixed modification. MS2 peak areas of each peptide were used for quantification.

Raw files from TMT-labeled samples were processed using Proteome Discoverer (version 2.5, Thermo Fisher Scientific) with the Sequest HT search engine. The same UniProt human proteome reference (20,384 entries) was used for database searching. Search parameters included: trypsin as the digestion enzyme with up to two missed cleavages; fixed modifications: carbamidomethylation of cysteine residues and TMTpro at peptide N-termini and lysine residues; variable modifications: methionine oxidation and protein N-terminal acetylation. The precursor mass tolerance was set to 10 ppm and fragment mass tolerance to 0.02 Da. The false discovery rate (FDR) was set to 1% at both the peptide and protein levels using the Percolator node. For quantification, TMTpro reporter ion intensities were extracted from MS2 spectra with the SPS (synchronous precursor selection) mass matching tolerance set to 20 ppm. Only unique peptides were used for protein quantification. Each channel was normalized to the total peptide amount within each multiplexed set.

Differential protein expression analysis was performed using the limma package (v3.62.1) in R. Proteins with adjusted P value < 0.05 (Benjamini–Hochberg correction) and |log₂(fold change)| > 0.5 were considered significantly differentially expressed. Volcano plots were generated using the ggplot2 package (v4.0.3) in R. Venn diagrams were generated using the VennDiagram package (v1.8.2). UpSet plots were generated using the UpSetR package (v1.4.1). Gene set enrichment analysis (GSEA) was performed using the clusterProfiler package (v4.8.3) in R.

## Statistical analysis

Statistical analyses were performed using GraphPad Prism 8 software. Unless otherwise specified, comparisons between groups were conducted using two-tailed Student’s t-tests. A p-value of less than 0.05 was considered statistically significant. *p < 0.05, **p < 0.01, ***p < 0.001.

---

**Extended Data Fig. 1.**
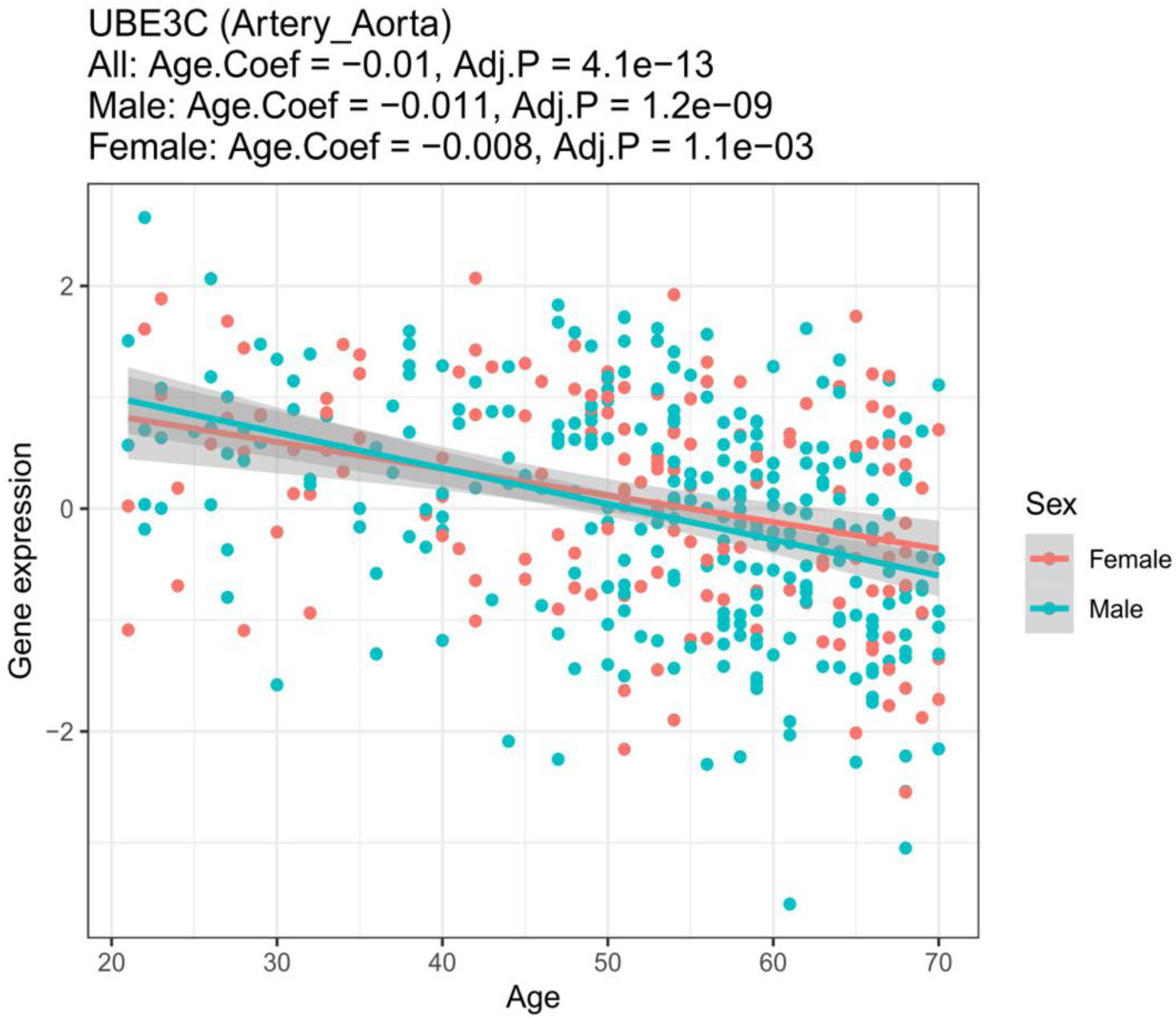
**Dot plot of UBE3C gene expression in human aorta according to GTEx analysis**. Each point represents an individual donor, with linear regression highlighting the significant correlation between UBE3C mRNA levels and age. Age.Coef and adj.P were indicated in male, female, and all sex.

**Extended Data Fig. 2.**
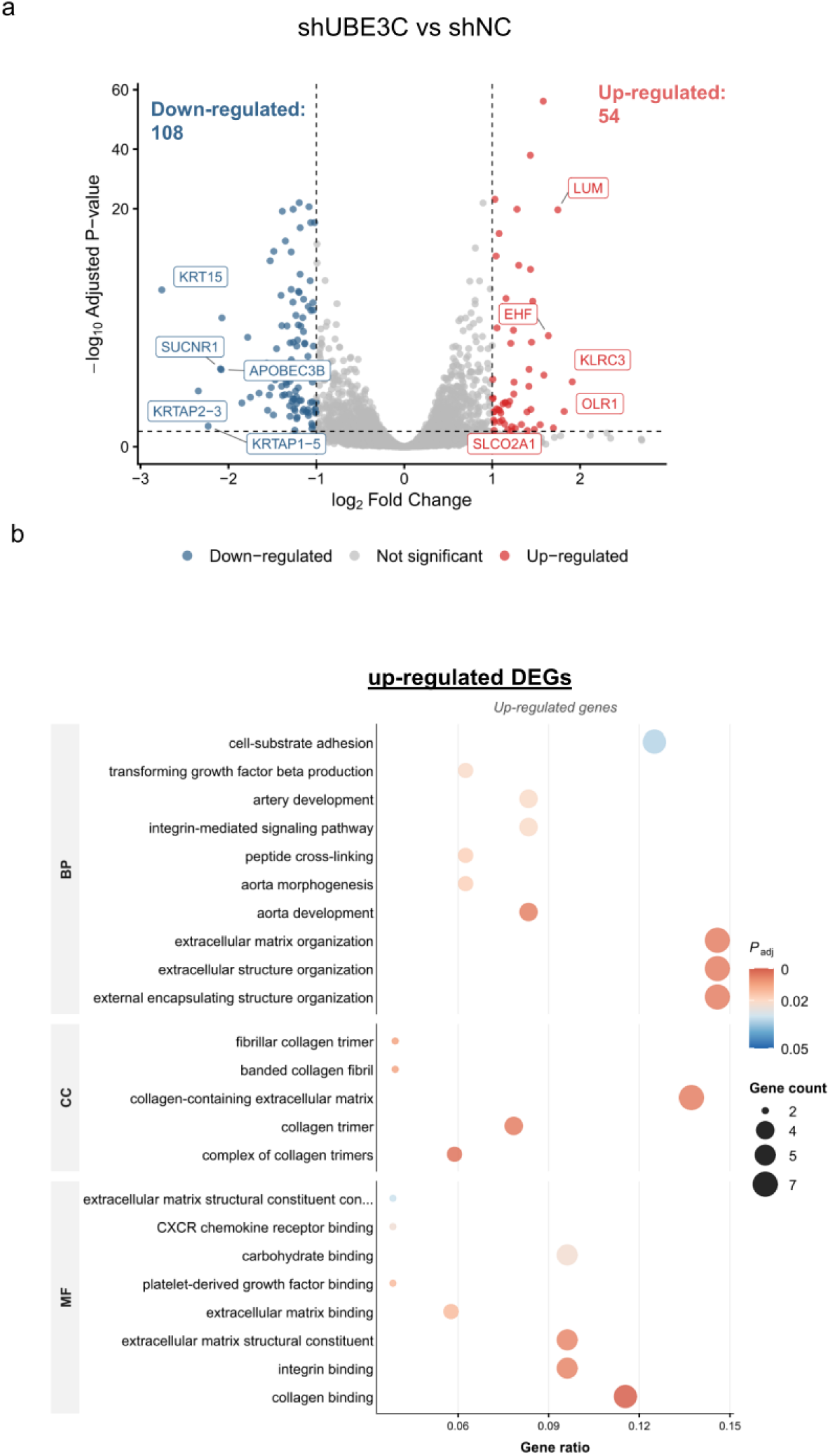
RNA-seq analysis of UBE3C-depleted hMSCs. a, Volcano plot of differentially expressed genes (DEGs) in UBE3C-knockdown versus control hMSCs (adj.P < 0.05, |log₂(fold change)| > 1). Red dots: significantly upregulated genes; blue dots: significantly downregulated genes; gray dots: non-significant genes. Top 5 upregulated and downregulated DEGs are annotated. **b**, Gene Ontology (GO) enrichment analysis of upregulated DEGs (adj.P < 0.05) in hMSCs at passage 1 (day 4 post-transduction), covering biological process (BP), cellular component (CC), and molecular function (MF) categories. The top 10 pathways were shown. n=2 biologically independent experiments.

**Extended Data Fig. 3.**
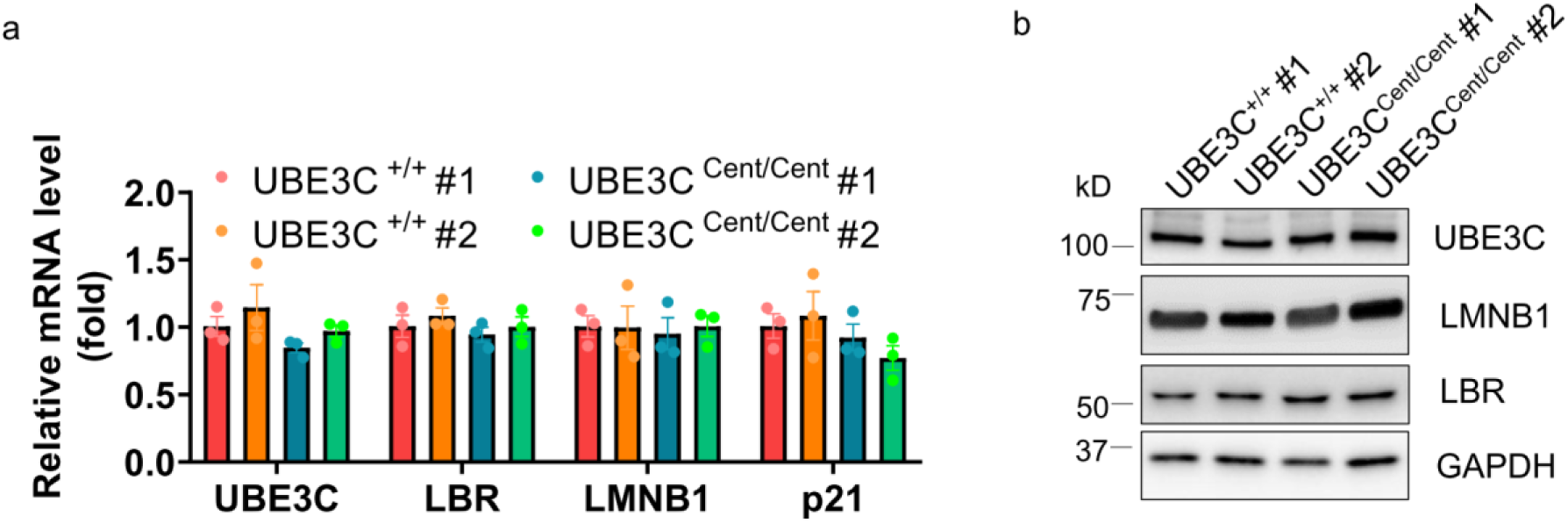
Expression of UBE3C, LMNB1 and LBR in UBE3C-KI hESCs. **a**, qRT-PCR analysis of *UBE3C*, *LBR*, *LMNB1*, and *CDKN1A* (p21) mRNA level in wild-type (UBE3C*^+/+^*#1 and #2) and longevity-associated UBE3C variant (UBE3C*^cent/cent^* #1 and #2) hESC clones. Data normalized to GAPDH (*n* = 3). **b**, Western blot analysis of UBE3C, LMNB1 (lamin B1), and LBR (lamin B1 receptor) protein expression in hESC clones as described in **a**. GAPDH served as a loading control. n=3 biologically independent experiments. Error bars represent mean ± SEM. *p < 0.05, **p < 0.01, ***p < 0.001 (unpaired Student’s t-test).

**Extended Data Fig. 4.**
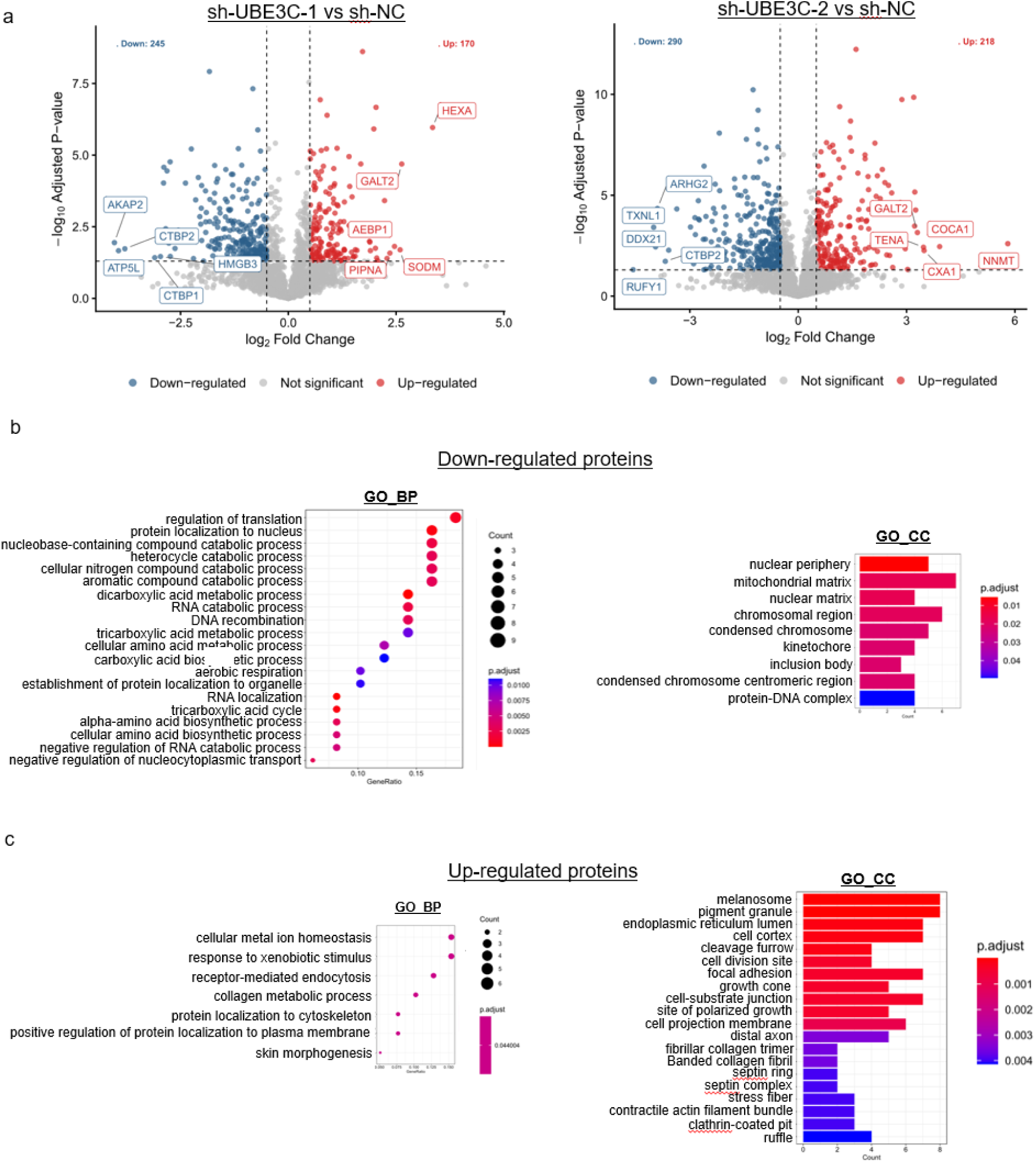
Proteomic alterations in UBE3C-depleted hMSCs. a, Volcano plot of differentially expressed proteins (DEPs) in hMSCs transduced with sh- UBE3C-1 or sh-UBE3C-2 versus sh-NC control (adj.P < 0.05, |log₂(fold change)| > 0.5). Red dots: significantly upregulated proteins; blue dots: significantly downregulated proteins; gray dots: non-significant proteins. Top 5 upregulated and downregulated DEPs are annotated. **b**, Gene Ontology (GO) enrichment of biological process (BP, left) and Cellular Component (CC, right) for overlapping down-regulated DEPs of sh-UBE3C-1 and sh-UBE3C-2 as described in **a**. **c**, GO enrichment of BP (left) and CC (right) for overlapping up-regulated DEPs. Top 20 significantly enriched pathways per category are shown (adj.P < 0.05).

**Extended Data Fig. 5.**
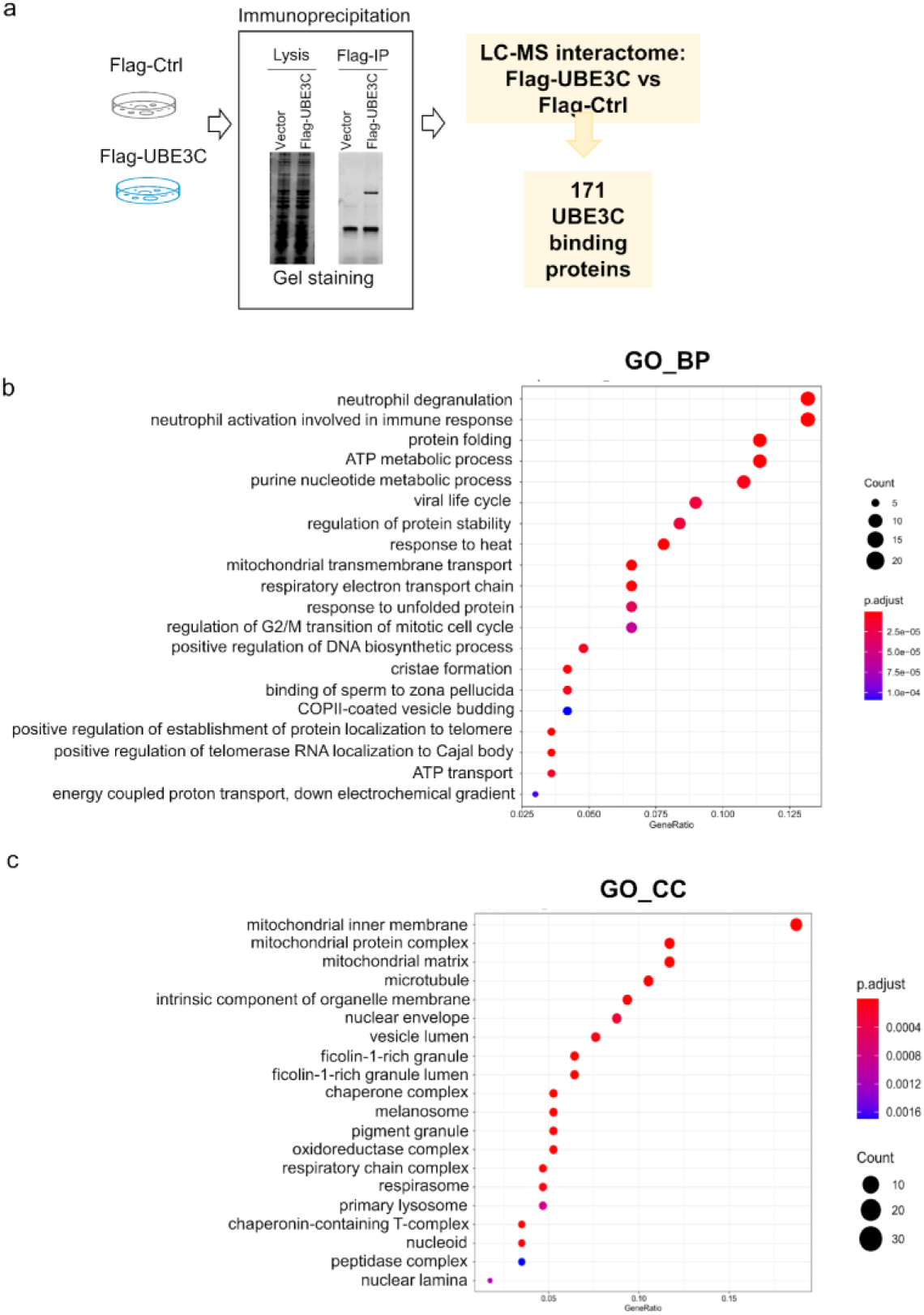
UBE3C interactome profiling by IP-MS. a, Experimental workflow for Flag-UBE3C immunoprecipitation coupled with mass spectrometry (IP-MS) in HEK293T cells. A total of 171 significant UBE3C-binding proteins were identified by comparing Flag-UBE3C interactome with Flag controls. n=3 biologically independent experiments. Adj.p < 0.05 **b**, Gene Ontology (GO) enrichment of biological process (BP) for UBE3C-interacting proteins (top 20 pathways, adj.P < 0.05) as described in **a**. **c**, GO cellular component enrichment of UBE3C-interacting proteins (top 20 pathways, adj.P < 0.05).

**Extended Data Fig. 6.**
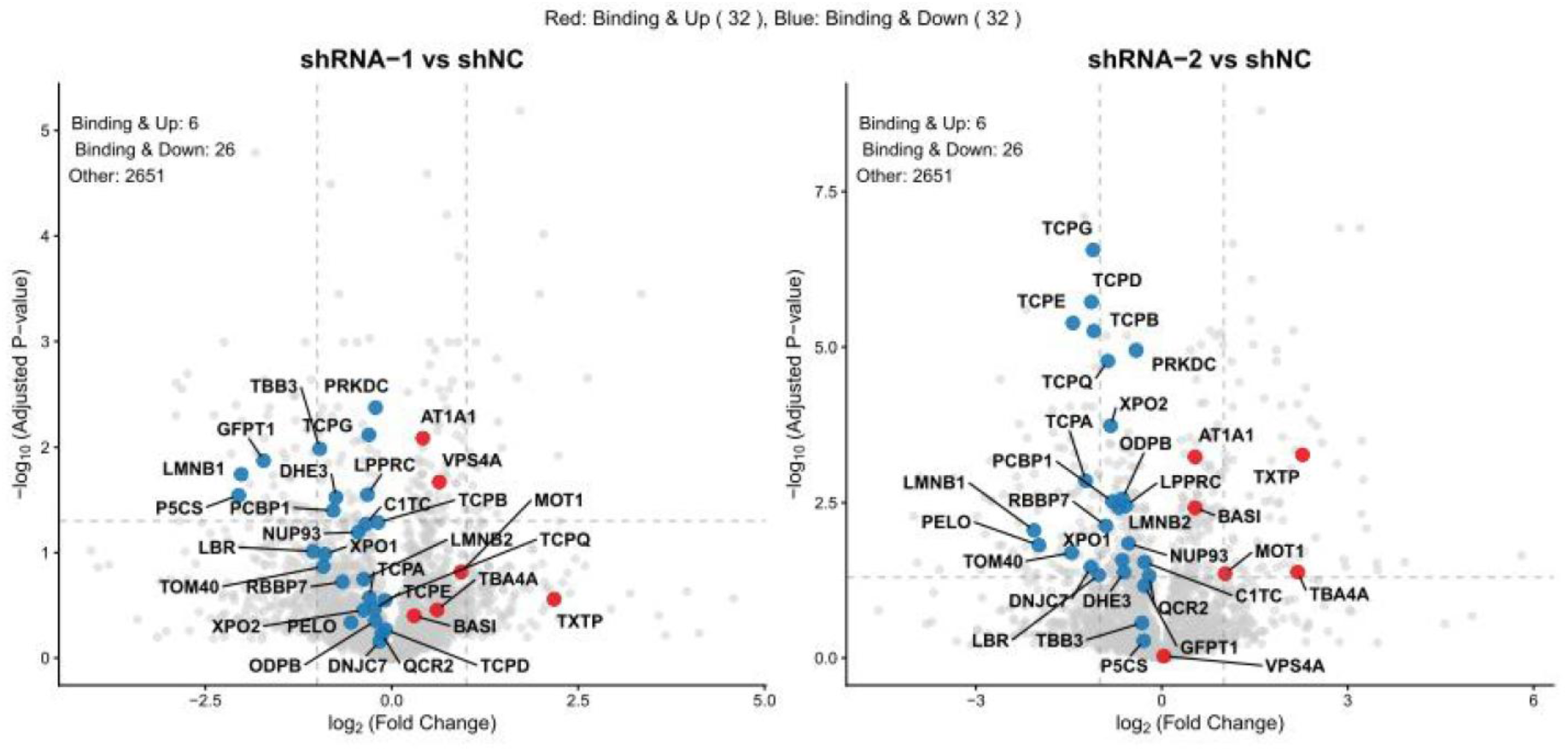
Identification of UBE3C binding proteins with expression levels affected by UBE3C depletion. Integrative analysis of the UBE3C interactome and differentially expressed proteins (DEPs) in hMSCs transduced with two independent shRNAs targeting UBE3C (shRNA-1, shRNA-2) versus shNC control. Upregulated (red) and downregulated (blue) DEPs overlapping with UBE3C binding proteins are annotated in the volcano plot. adj.P < 0.05, |log₂(fold change)| > 0.5.

**Extended Data Fig. 7.**
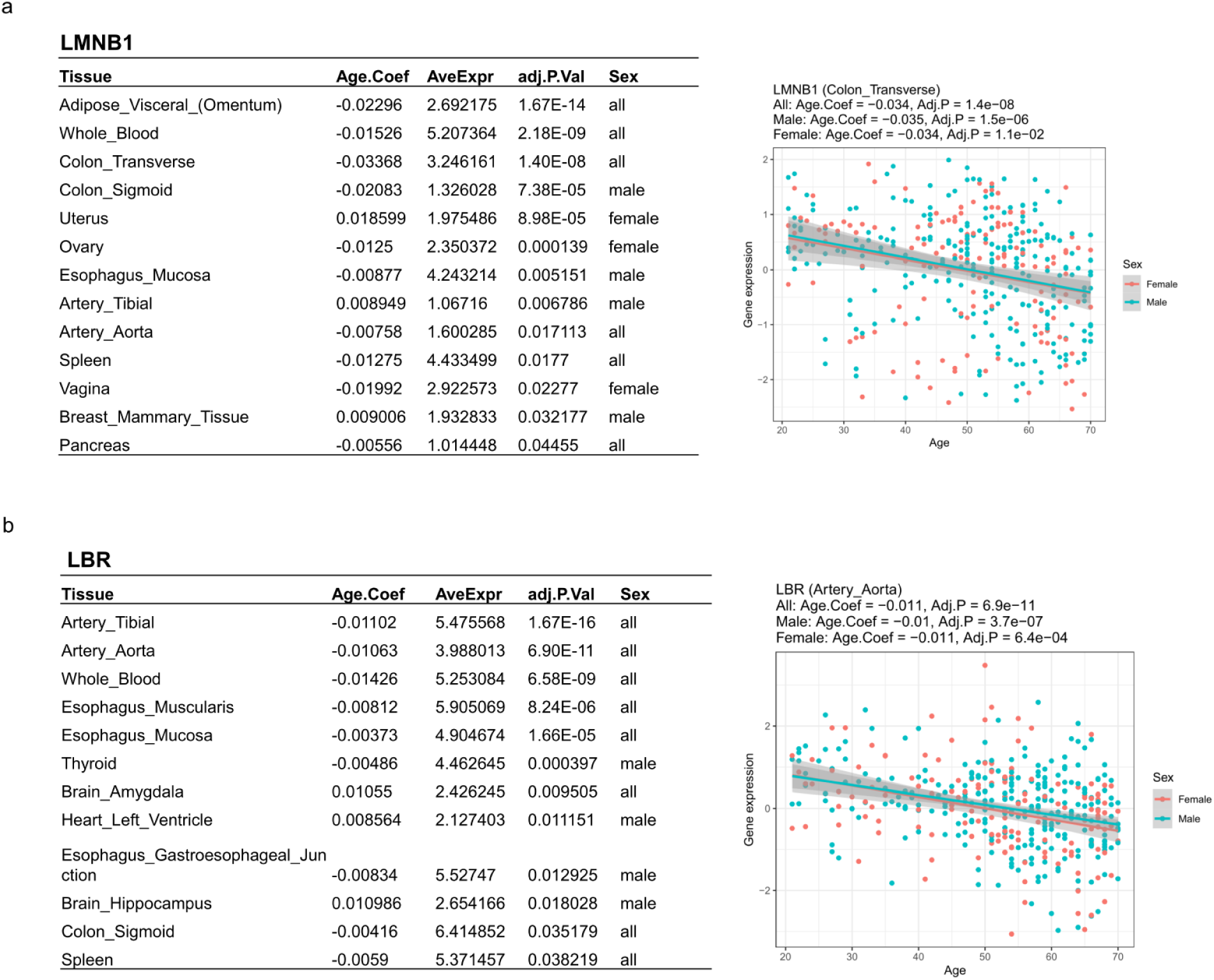
Tissue-specific decline of nuclear lamina components LMNB1 and LBR during human aging. a, Age-associated changes in LMNB1 expression. Left: Linear regression analysis across GTEx tissues showing significant negative correlations with donor age. Negative values of Age. Coef indicate expression decline with aging. Right: Representative LMNB1 mRNA levels with age in transverse colon. Each dot represents an individual donor. Age range is indicated. **b**, Age-associated changes in LBR expression. Left: Regression analysis across GTEx tissues. Right: LBR mRNA level in aortic artery.

**Extended Data Fig. 8.**
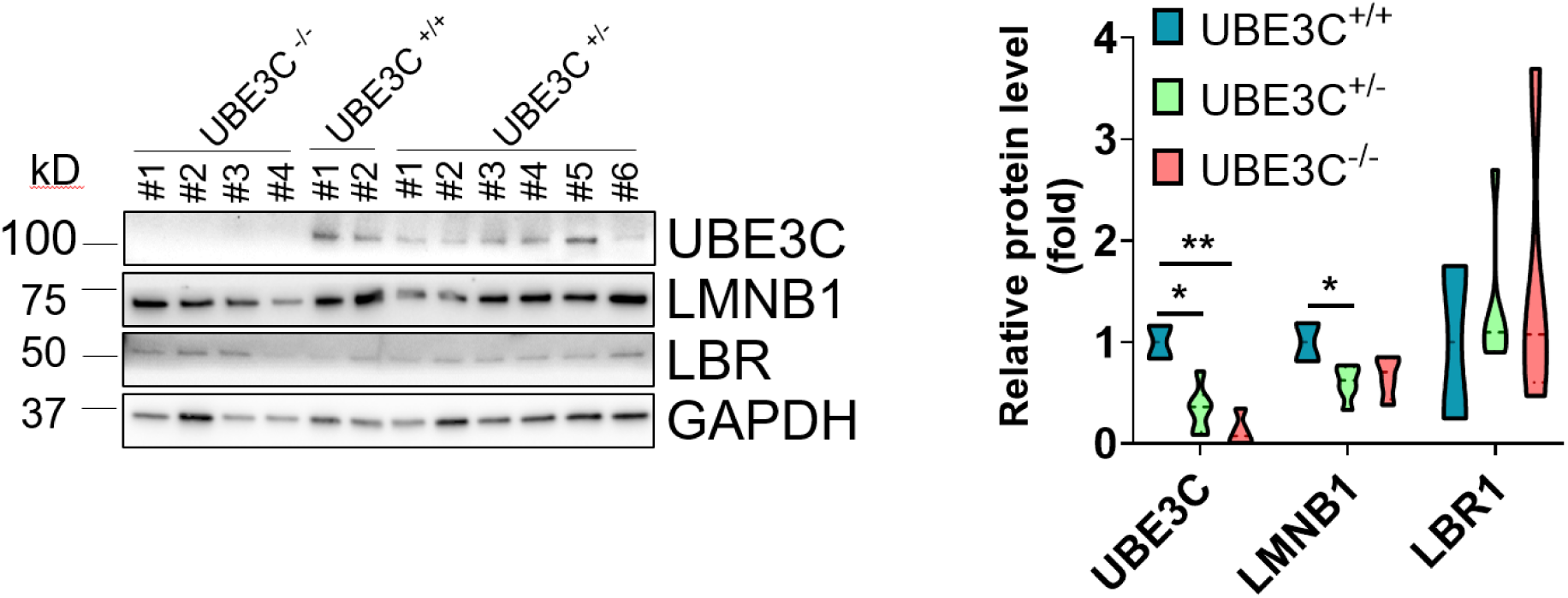
UBE3C knockdown leads to reduced LMNB1 protein levels in mice. Left panel: Western blot of UBE3C, LMNB1, and LBR in ear tissues from UBE3C*^-/-^* (*n*=4), UBE3C*^+/-^*(*n*=6), and UBE3C*^+/+^* mice (*n*=2) mice. Right panel: Quantification of UBE3C, LMNB1, and LBR protein levels normalized to GAPDH. Error bars represent mean ± SEM. *p < 0.05, **p < 0.01, ***p < 0.001 (unpaired t-test unless noted).

**Extended Data Fig. 9.**
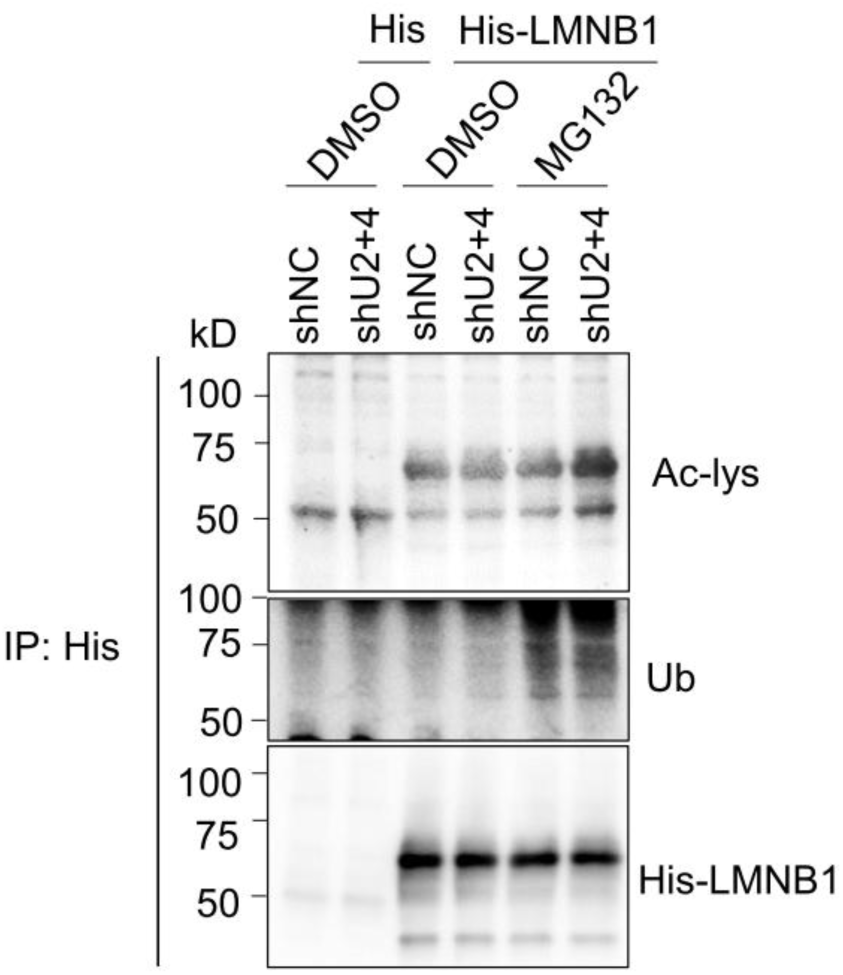
**Ubiquitination level of LMNB1 is not changed by UBE3C depletion.**Detection of ubiquitination and acetylation modifications on His-tagged LMNB1 in HEK293T cells following UBE3C knockdown. Cells were transfected with sh-UBE3C or sh-NC, treated with 10 μM MG132 (proteasome inhibitor) or DMSO (6 hr). Immunoblotting shows acetylation (Ac-lys) and ubiquitination (Ub) levels of His-LMNB1.

**Extended Data Fig. 10.**
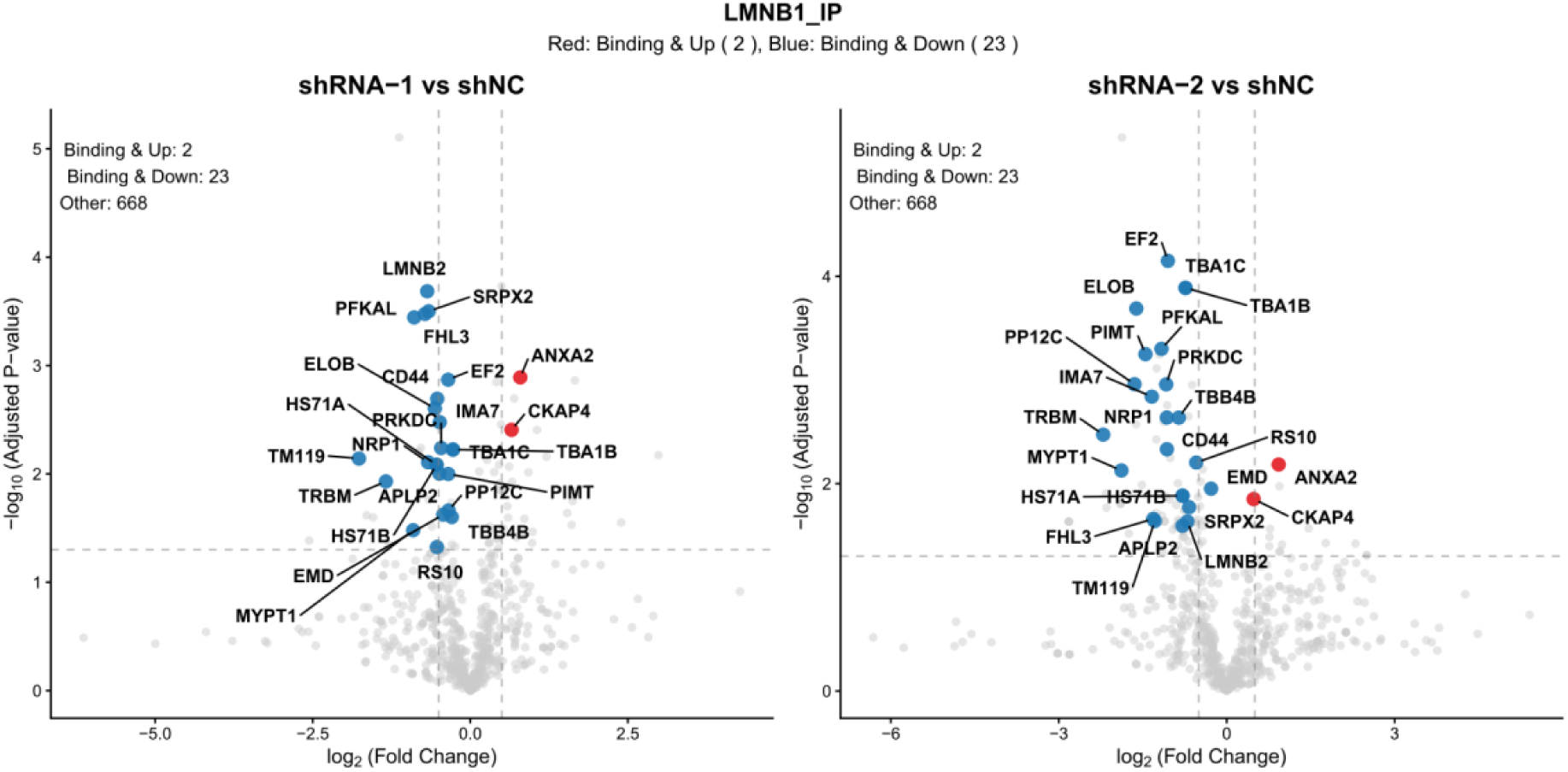
Identification of LMNB1-interacting proteins with binding affected by UBE3C depletion. Volcano plot showing differentially binding proteins (DBPs) in hMSCs transduced with two independent shRNAs targeting UBE3C (shRNA-1, shRNA-2) versus shNC control. Upregulated (red) and downregulated (blue) DBPs of LMNB1 are annotated (adj.P < 0.05, |log₂(fold change)| > 0); non-significant DBPs are in gray. n=biologically independent experiments.

**Extended Data Fig. 11.**
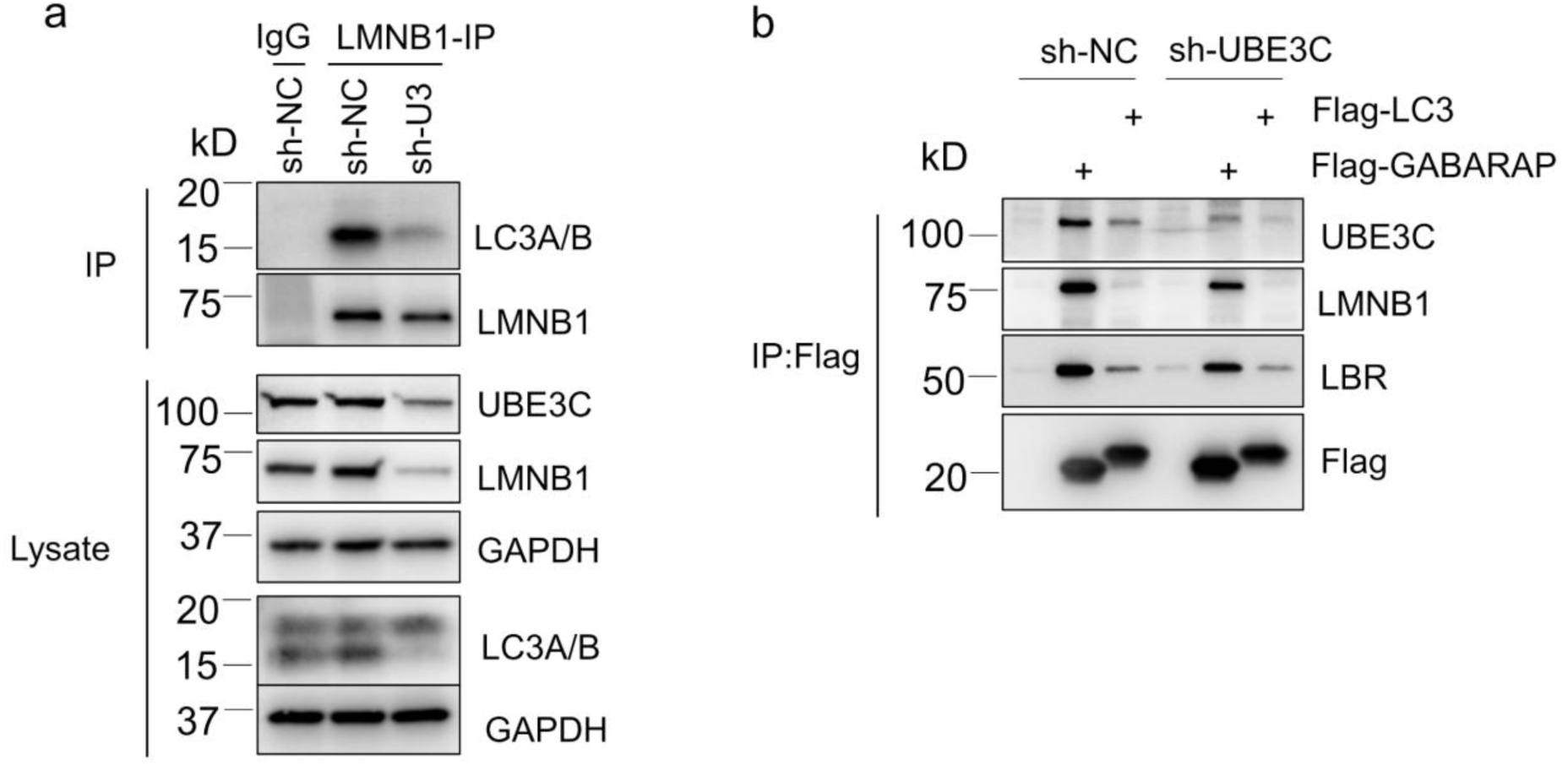
UBE3C depletion reduces LMNB1/LBR interactions with LC3/GABARAP. a, Endogenous co-immunoprecipitation (Co-IP) of LC3A/B with LMNB1 in human mesenchymal stromal cells (hMSCs) following UBE3C knockdown. hMSCs were transduced with lentiviral shRNA targeting UBE3C (sh-UBE3C) or non-targeting control (sh-NC), followed by anti-LMNB1 IP and immunoblotting for LC3A/B. **b**, Co-IP analysis of UBE3C, LMNB1, and LBR interactions with overexpressed Flag-LC3 or Flag-GABARAP in HEK293T cells. Cells were transfected with sh-UBE3C or sh-NC prior to Flag-IP and immunoblotting for the indicated proteins.

